# A bile metabolite atlas reveals infection-triggered interorgan mediators of intestinal homeostasis and defense

**DOI:** 10.1101/2023.03.04.531105

**Authors:** Ting Zhang, Yuko Hasegawa, Matthew K. Waldor

## Abstract

An essential function of the liver is the formation of bile. This aqueous solution is critical for fat absorption and is transported to the duodenum via the common bile duct. Despite extensive studies of bile salts, other components of bile are less well-charted. Here, we characterized the murine bile metabolome and investigated how the microbiota and enteric infection influence bile composition. We discovered that the bile metabolome is not only substantially more complex than appreciated but is dynamic and responsive to the microbiota and enteric infection. Hepatic transcriptomics identified enteric infection-triggered pathways that likely underlie bile remodeling. Enteric infections stimulated elevation of four dicarboxylates in bile that modulated intestinal gut epithelial and microbiota composition, inflammasome activation, and host defense. Our data suggest that enteric infection-associated signals are relayed between the intestine and liver and induce transcriptional programs that shape the bile metabolome, modifying bile’s immunomodulatory and host defense functions.

## Introduction

The cross-kingdom interplay between microbiomes and their mammalian hosts generates a diverse pool of compounds, many of which enter the circulatory system and impact host organ function^1–3^. The liver receives blood from the portal vein, which drains the intestine, and from the systemic circulation, allowing it to play a critical role in integrating chemical signals from the diet and the gut microbiome with those present in systemic blood (Fig. 1a). The liver responds to these microbial stimuli by synthesizing a large array of compounds, including innate immune effectors, hormones, and macronutrients (e.g., vitamins). These compounds are secreted into the systemic circulation via the hepatic vein and modulate host physiology. In addition, infection or tissue injury can trigger a hepatic response referred to as the acute phase response, in which hepatocytes produce and secrete large amounts of proteinaceous mediators, such as complement proteins, serum amyloid A, and hemopexin into systemic circulation that facilitate host defense and /or repair at distal sites^4, 5^.

**Figure 1.**
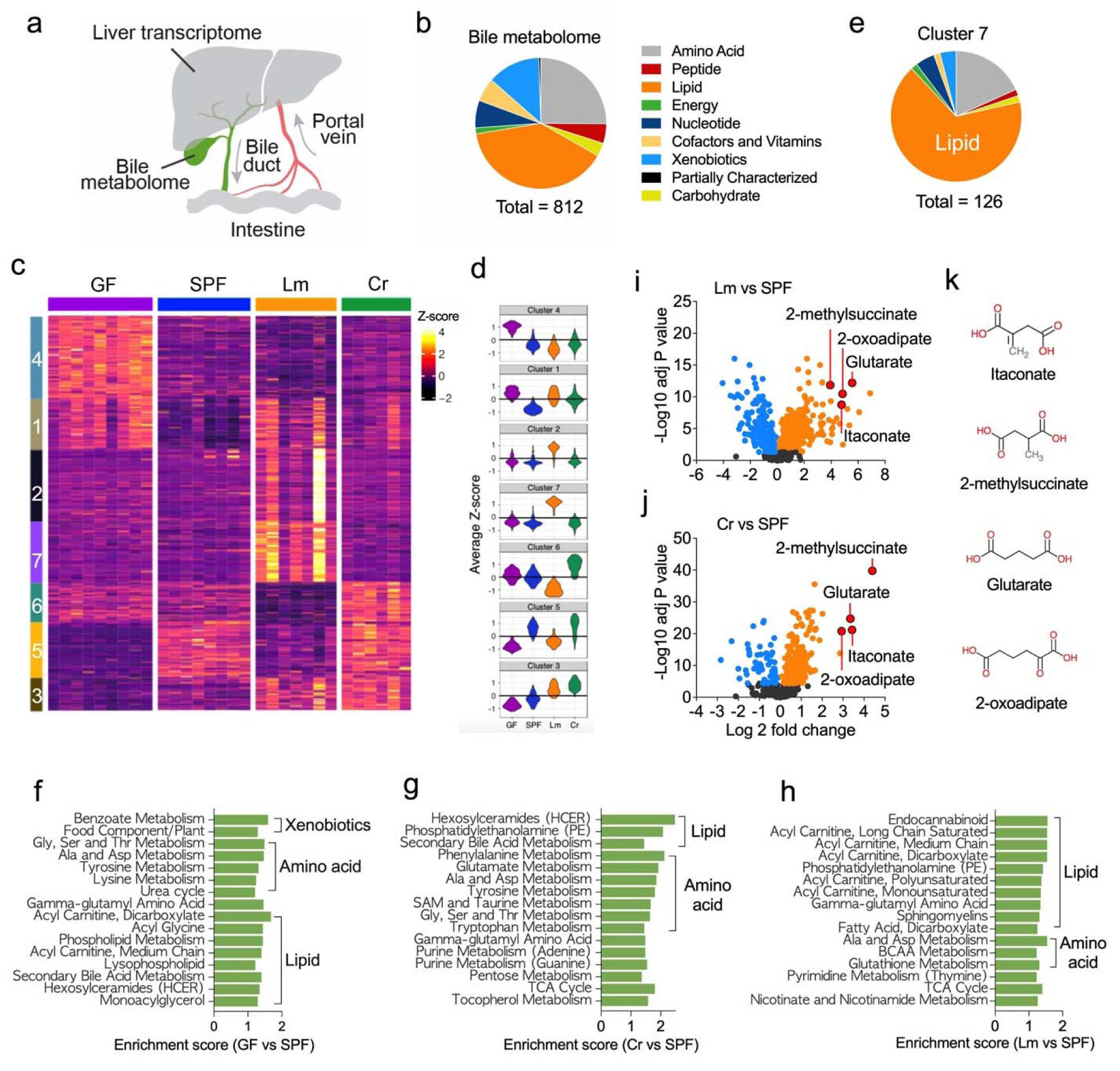
The microbiota and enteric infection modify the bile metabolome. (a) Schematic of enterohepatic circulation. The liver receives blood from portal vein that drains the intestine and produces bile that is delivered to the duodenum via the common bile duct. (b) Functional categories of the 812 bile metabolites in SPF mice that were identified by global metabolomic profiling. (c) Unsupervised clustering (k-medoids) of the bile metabolites identified in four groups of mice (germ free (GF), specific pathogen free (SPF), *Listeria monocytogenes* infected (Lm), and *Citrobacter rodentium* (Cr) infected) based on their patterns of relative abundance. Each row represents a metabolite and each column represents one sample. (d) Violin plot of the distribution of the Z-scores of the metabolites in each cluster. (e) Functional categories of bile metabolites in cluster 7 identified by unsupervised clustering. (f-h) Enriched pathways of metabolites with differential abundance between SPF and GF mice (f), SPF and Cr mice (g), and SPF and Lm mice (h). (i-j) Differential abundance of bile metabolites in *L. monocytogenes* (i) or *C. rodentium* (j) infected animals compared to uninfected mice. Blue, black and orange dots represent metabolites with reduced, unchanged, or increased abundance in the infected samples. (k) Structural formulas of infection-stimulated bile dicarboxylates.

Besides synthesizing many of the non-cellular components of blood, the liver also plays a central role in formation of bile, an aqueous solution that can be stored in the gallbladder prior to its transport to duodenum via the common bile duct (Fig 1a)^6^. Bile consists of lipids, proteins, metabolites and bile acids ^7^. Bile acids are a principal component of bile and play an essential role in lipid absorption; furthermore, these sterols regulate host metabolism^8^, antimicrobial defense^9^ and differentiation of intestinal immune cells ^10, 11^. Primary bile acids are synthesized in the liver, converted to secondary bile acids by the gut microbiota and are re-absorbed by the portal circulation back into the liver as part of a circuit referred to as enterohepatic circulation. The liver monitors the composition of bile acids in the portal and systemic blood to regulate *de novo* synthesis of primary bile acids^12^. Although the functions of bile acids have received considerable attention ^13–15^, the chemical composition, functions, and regulation of other bile constituents has not been the subject of much experimental scrutiny. In contrast to the widely surveyed serum metabolome^1, 3^, changes in bile composition in response to the microbiota or to enteric infection are largely unknown.

Here, we used global metabolomic analyses to characterize the murine bile and to investigate how the composition of bile is altered by the microbiota and enteric infection. We identified >800 bile metabolites, many of which were not known to be bile components. There were marked differences in bile metabolites in the absence of the microbiota. Moreover, infection with pathogens that vary in their dissemination from the intestine were found to modify the bile metabolome in both shared and distinctive fashions. We also discovered that bile dicarboxylates that were elevated by enteric infections modulate gut epithelial and microbiota composition, intestinal inflammasome activity, and host defense against an enteric pathogen. Collectively, our findings reveal that bile composition is highly complex, responsive to the microbiota and infection, and functions in an interorgan innate defense circuit that links the liver and intestine.

## Results

### The bile metabolome is complex

We used global metabolomic profiling to create an atlas of murine bile metabolites. Bile samples were obtained from the gallbladders of C57BL/6 specific pathogen free (SPF) animals. In total, 812 metabolites, representing 9 functional categories, were identified in the bile (Fig. 1b, Table S1). Many of the compounds identified were not known as bile components. For example, several monoacylglyercols were found among bile lipids, the dominant class of bile components (Fig 1b, Fig 2). Bile metabolites were not only of host origin. Many of the compounds in the bile of SPF mice are generated or processed by the gut microbiota and have been identified in the host portal and systemic circulation (e.g., equol sulfate and indoxyl sulfate)^16, 17^ (Fig. S1a, S2c). Furthermore, several bile constituents, such as benzoate, hydroxycinnamate, and genistein, are of dietary origin (Fig. S1a). The presence of microbial-and dietary-derived compounds in bile suggests that they are distributed thorough enterohepatic circulation and following their intestinal absorption, these compounds are processed by the liver and secreted in bile, in a similar fashion as described for xenobiotics ^16^.

**Figure 2.**
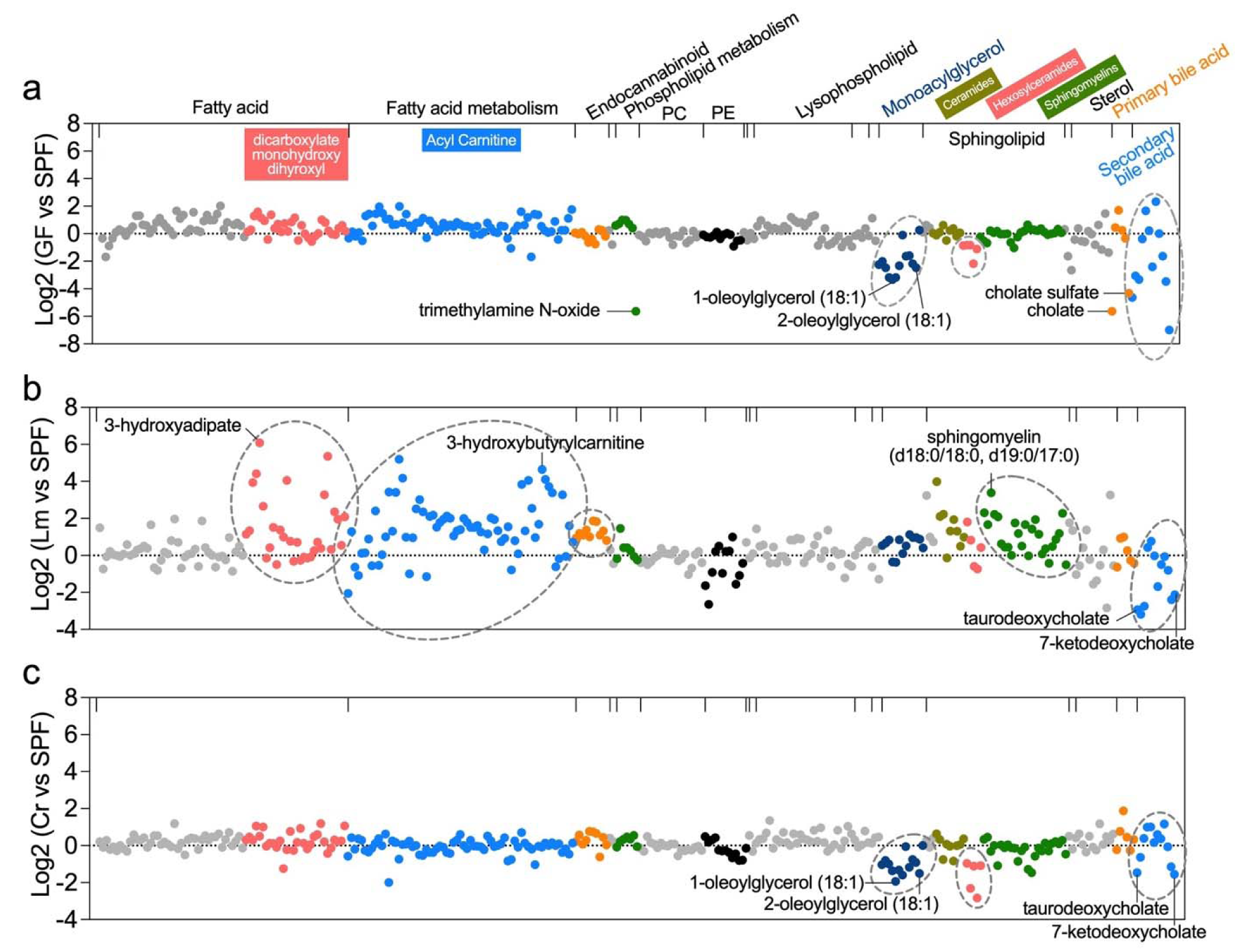
The bile lipidome is shaped by the microbiota and modified by enteric infection. (a-c) Differential abundance of bile lipids in germ free (GF) (a), *L. monocytogenes* (Lm) infected (b), or *C. rodentium* (Cr) infected mice (c) compared to SPF animals. The categories of lipids are labeled on the top of the graph in (a). Differentially abundant lipids that were identified by pathway enrichment analysis are labeled with dished circles.

To investigate how the microbiota and enteric infection impacts bile composition, we compared the metabolomes of bile from SPF mice to that from C57BL/6 germ-free (GF), and mice orally infected with *Listeria monocytogenes* (Lm) or *Citrobacter rodentium* (Cr). An unsupervised clustering algorithm (k-medoids) was used to classify the bile metabolites identified in the 4 conditions based on their patterns of abundance. The 812 bile metabolites were partitioned into 7 clusters, which in aggregate distinguished the conditions (Fig. 1c, d, Table S1). Principal component analysis (PCA) and pathway enrichment analysis also revealed that the bile metabolomes in these 4 conditions were distinct (Fig. 1f-h, S1b).

### Bile metabolites are shaped by the microbiota

The makeup of bile in SPF and GF mice was easily distinguishable (clusters 4, 1, 5, and 3, Fig. 1c, d). The relative abundance (as reflected in signal intensities) of nearly 60% of the metabolites differed between the two groups (p<0.05, Fig. S1c). As expected, many compounds that are generated or processed by the gut microbiota (e.g., N-acetylhistamine and indolepropionylglycine) were found in the bile of SPF and not GF animals (Fig. S2c). Compounds classified as xenobiotics, amino acids, and lipids were the most affected by the absence of the microbiota (Fig. 1f).

The bile lipidome was distinct in GF mice. As expected, known microbial-derived lipids such as secondary bile acids, short chain fatty acids (e.g., butyrate), and trimethylamine N-oxide had reduced abundance in GF vs SPF animals (Fig. 2a). The absence of the microbiota was also associated with reduced abundance in bile of several monoacylglycerols (e.g., 1-oleoylglycerol (18:1) and 2-oleoylglycerol (18:1)) and hexosylceramide (Fig. 2a), suggesting that the microbiota contributes to the production or metabolism of these lipids. Together, these observations suggest that the interplay between the microbiota and host has a profound impact on bile composition.

### Enteric infection modifies bile composition

Profiling of bile metabolites was also carried out four and ten days following oral inoculation of mice with *L. monocytogenes* and *C. rodentium* respectively. In this model, *L. monocytogenes* routinely disseminates from the intestine to the liver and gallbladder following oral inoculation^18^. In contrast, *C. rodentium*, a natural murine enteric pathogen, primarily replicates in the cecum and colon. At four days post infection (dpi), the burden of *L. monocytogenes* in the liver and the gallbladder peaks ^19^; in the latter organ, the pathogen replicates extracellularly in bile where it reaches concentrations of ∼10^9^ CFU/ml (Fig. S1f, g). *C. rodentium* was never isolated from the gallbladder and a peak pathogen burden of ∼10^9^ CFU was observed in the colon 10 dpi (Fig. S1h). At this point, in some animals, a small number (median, 400) of *C. rodentium* CFU were isolated from the liver (Fig. S1i). Both pathogens altered bile metabolite profiles compared to SPF animals, though the clusters of changing metabolites differed between pathogens (Fig. 1c, d, g, h; S1c, d). Thus, even in the absence of pathogen invasion of the gallbladder, enteric infection leads to remodeling of the bile metabolome in a pathogen-specific manner.

*L. monocytogenes* infection altered the abundance of more bile metabolites (529) than *C. rodentium* infection (332, Fig. 1c, S1d, e). Metabolites in cluster 2 and 7, which were highly enriched for lipids (Fig. 1e), were only heightened in *L. monocytogenes* infected animals (Fig. 1c, d). Pathway enrichment analysis of all differentially abundant metabolites also revealed that *L. monocytogenes* infection triggered alteration of the bile lipidome (Fig. 1h, 2b, S1d). In contrast to the GF, SPF and Cr groups of animals, bile from the Lm group had elevated abundances of fatty acids, acyl-carnitines, endocannabinoids, and sphingomyelins (Fig. 2b), potentially in part due to the activities of the pathogen’s phospholipases^20^. In contrast, *C. rodentium* infected animals, like GF mice, had decreased abundance of hexosylceramides and monoacylglycerols (Fig. 2a, c), suggesting that alterations in the microbiota caused by the proliferation of *C. rodentium* in the intestine ^21^ reduced the abundance of these likely microbiota-derived bile lipids.

Bile composition in the Lm and Cr groups also exhibited some similarities. For instance, the abundance of several secondary bile acids was reduced in both infections (e.g., taurodeoxycholate and 7-ketodeoxycholate, Fig. 2b, c,), suggesting that enteric infection disrupts the production and/or enterohepatic circulation of these microbiota-derived lipids as observed in Gautam et. al. ^22^. The abundances of metabolites in cluster 3 were elevated in both the Lm and Cr groups (Fig. 1c, d). In total, the amounts of 138 bile metabolites were elevated by these two enteric pathogens relative to the SPF group (Fig. S2a, Table S2). The common enteric infection induced metabolites were enriched in the energy and amino acid categories (Fig. S2b, d, e, g, h). Increased abundances of four dicarboxylates (Fig. 1i, j, k), 2-methylsuccinate, glutarate, 2-oxoadipate, and itaconate, were particularly prominent in both infections (Fig.1 i, j, and S2d, e). Targeted LC-MS/MS analysis confirmed the elevation in the abundance of these 4 dicarboxylates in individual *C. rodentium* infected animals (Fig. S2i). The former three compounds are intermediate products of amino acid metabolism (Fig. S2d, e), whereas itaconate is generated from the TCA cycle metabolite aconitate (Fig. S2d, e). Thus, enteric infection with *C. rodentium* and *L. monocytogenes* trigger shared and distinct modifications in the composition of bile. Together, these data reveal that the bile metabolome is not only highly complex, but markedly influenced by the microbiota and altered in distinct ways by enteric infections. Below, we investigate potential pathways that generate these compounds and phenotypes linked to some of these dicarboxylates, none of which were previously known to be bile components.

### Enteric infections alter the hepatic transcriptome

Given the central role of the liver in bile formation, we profiled the hepatic transcriptome at the same time points as we assayed bile metabolites to investigate if changes in hepatic gene expression patterns underlie some of the changes observed in the bile metabolome during *L. monocytogenes* and *C. rodentium* infection. Transcriptomes from *C. rodentium* infected animals were divided into two groups based on whether or not they had detectable (Cr positive) or undetectable (Cr negative) *C. rodentium* in their livers at the time of sacrifice and analyzed separately. In total, 3877 hepatic transcripts had altered abundance in response to *L. monocytogenes* infection (2160 with increased abundance and 1717 with decreased abundance). In *C. rodentium* infected animals, there were fewer differentially expressed genes (Cr positive: 2414 increased and 900 decreased, Cr negative: 1113 increased and 307 decreased, Table S3). Principal component analysis clearly separated the transcriptional profiles from the 4 conditions (Fig. S3a). Importantly, the transcriptional profiles observed in the Cr negative group of animals were distinct from those found in SPF mice (Fig. S3a), consistent with the idea that enteric infection modulates hepatic transcriptional programs even in the absence of detectable pathogen dissemination to the liver.

Unsupervised clustering was used to partition differentially expressed genes (DEGs) into 7 clusters (Fig. 3a, b, adjusted p-value < 0.05). This analysis revealed that there were common changes in hepatic transcriptome profiles associated with infection (Fig. 3a, b). Gene set enrichment analysis and gene-metabolite network analysis also revealed that *L. monocytogenes* and *C. rodentium* infection affected similar biological processes (Fig. 3c, Fig. S4). Notably, both enteric infections stimulated hepatic expression of genes linked to inflammation (Fig. 3c), demonstrating that the liver mounts an inflammatory response even in the absence of pathogen dissemination from the intestine. In contrast, expression of many genes related to metabolism, including amino acid metabolism (KEGG 00260 in Fig. 3d, S4) and detoxification (KEGG 00980 in Fig. 3d) were markedly reduced in both infection models, suggesting that enteric infection depresses a subset of hepatic functional programs. In particular, infection reduced expression of liver genes involved in oxidative phosphorylation, including all four enzyme complexes in the electron transport chain (ETC) (Fig. S5a-d). Similarly, the abundance of transcripts encoding several tricarboxylic acid (TCA) cycle enzymes (Fig. 4a), including Idh1 and Sdh complex components, was reduced (Fig. 4b, c). In contrast, the abundance of transcripts for several enzymes in the glycolysis pathway was increased (Fig. 3c, Fig. 4d). Together, these observations suggest that the liver switches from oxidative phosphorylation to glycolysis for energy production during enteric infection. A similar switch is observed during stimulation of many immune cells ^23, 24^, in part due to the activity of pro-inflammatory cytokines ^25^, suggesting that these cytokines may have a similar impact on hepatic metabolism.

**Figure 3.**
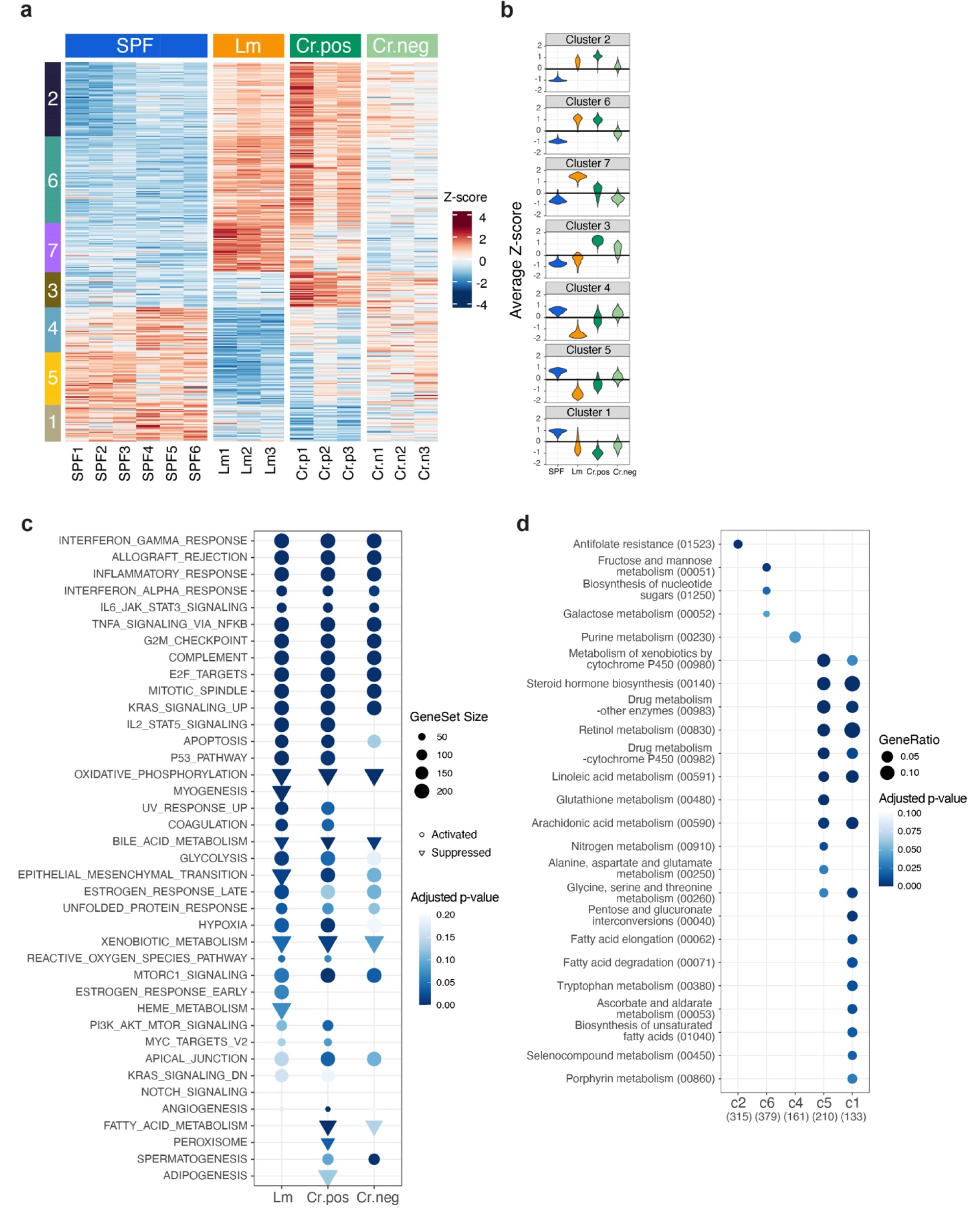
Hepatic transcriptome profiles are altered by enteric infection. (a) Unsupervised clustering (k-medoids) of the differentially expressed genes (DEGs) identified in the 4 conditions (uninfected (SPF), *Listeria monocytogenes* infected (Lm), and *Citrobacter rodentium* infected with detectable (Cr. pos) or undetectable (Cr. neg) CFU in the liver), based on their patterns of relative abundance. Each row represents a gene and each column represents one mouse. (b) Violin plot of the distribution of the Z-scores of the transcript abundance in each cluster. (c) Gene set enrichment analysis of differentially expressed genes (DEGs) that were identified in Lm vs SPF, Cr liver positive vs SPF, and Cr liver negative vs SPF conditions. (d) Functional enrichment analysis of differentially expressed genes using a subset of the KEGG database restricted to metabolism pathways. The categories on the x-axis correspond to the clusters in Fig. 3a.

**Figure 4.**
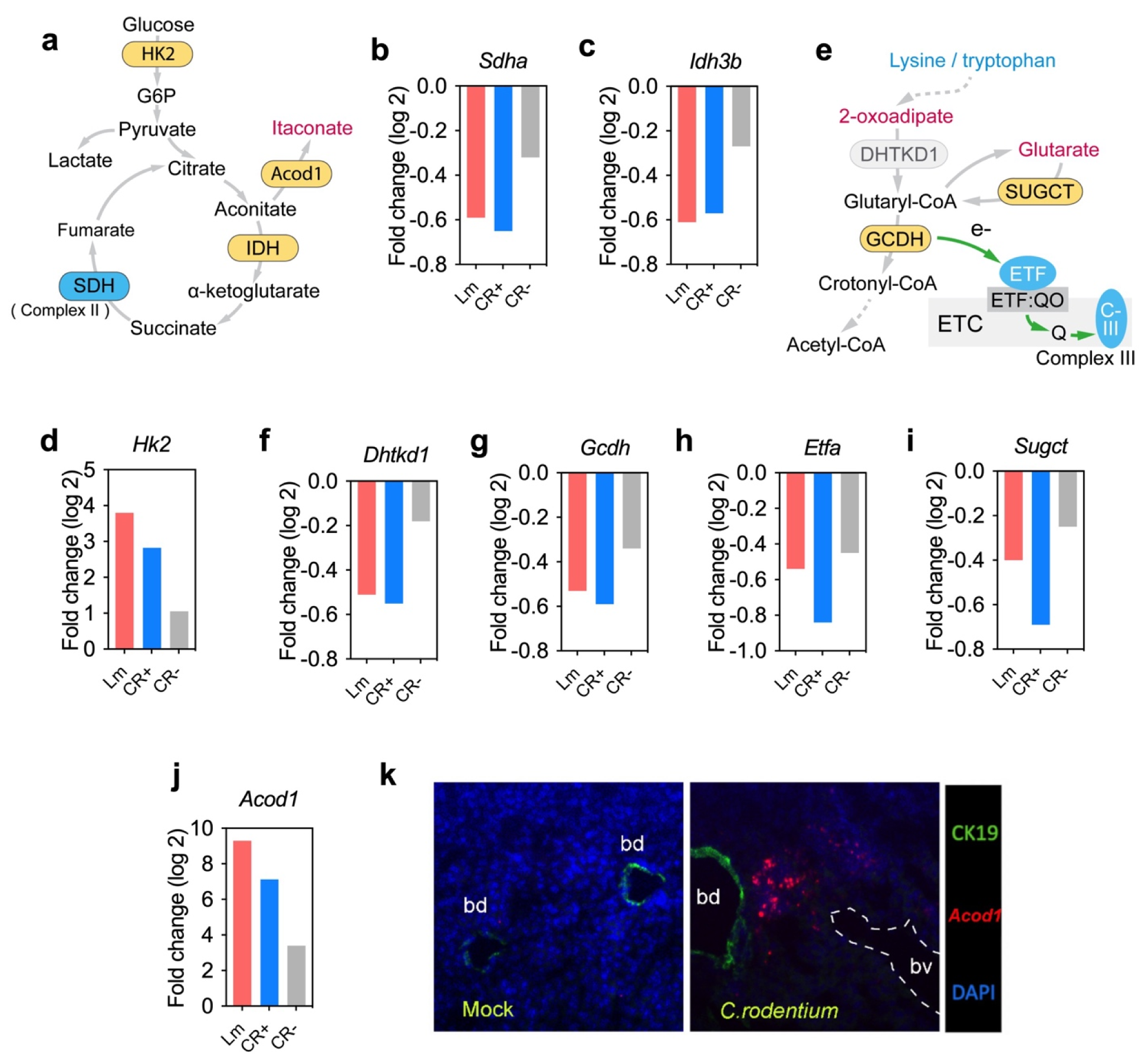
Hepatic gene expression alterations that contribute to infection-associated changes in the abundance of bile dicarboxylates. (a) Schematic of pathway for itaconate production. Sugars such as glucose are processed for energy production by Hexokinase 2 (HK2) and further converted to pyruvate, which in turn feeds the TCA cycle. Itaconate is produced from the TCA cycle metabolite aconitate by the enzyme Acod1. (b-d) The relative expression levels of TCA cycle enzymes *Sdha* (b) and *Idh3b* (c), glycolysis enzyme *Hk2* (d) in liver in enteric infection. (e) Proposed model for generation of glutarate and 2-oxoadipate as intermediate products of lysine and tryptophan metabolism. Mutations of GCDH lead to elevations of glutarate, whereas mutations of SUGCT and DHTKD1 lead to elevations of glutarate and 2-oxoadipate respectively ^28, 29^. GCDH is a FAD-dependent enzyme that requires the electron-transferring-flavoprotein (ETF) complex to transfer electrons to the electron transfer chain (ETC); Q, ubiquinone, ETF:QO, electron transfer flavoprotein-ubiquinone oxidoreductase. (f-i) The relative expression level of *Dhtkd1* (f), *Gcdh* (g), *Etfa* (h), and *Sugct* (i) in murine liver during enteric infection. (j) The relative expression levels of *Acod1* in liver in enteric infection. (k) RNA-FISH analysis of *Acod1*(red) in the liver; CK19 (green) bile duct marker, DAPI (blue), nuclei.

We next asked whether the hepatic transcriptome could explain the increases we observed in specific novel bile metabolites, in particular the four dicarboxylic acids that were sharply increased in bile samples from infected mice (Fig. 1i, j). Glutarate and 2-oxoadipate are intermediate products of lysine and tryptophan catabolism ^26^ (Fig. 4e), but the processes that govern their abundance are not fully characterized. Increased excretion of glutarate and 2-oxoadipate in urine is observed in glutarate aciduria type I (GA-1) and 2-oxoadipic aciduria (OA) ^27, 28^. GA-1 is caused by loss of function mutations in glutaryl-CoA dehydrogenase (GCDH), the enzyme that mediates glutaryl-CoA dissimilation ^29^, whereas OA is due to mutation in the 2-oxoadipate catabolism enzyme 2-oxoglutarate-dehydrogenase-complex-like protein (DHTKD1)^28^. The abundance of GCDH and DHTKD1 mRNA was reduced during *L. monocytogenes* and *C. rodentium* infection (Fig. 4f, g), likely explaining the corresponding elevated abundance of glutarate and 2-oxodipate in bile. Furthermore, GCDH is a FAD-dependent enzyme and its function relies on the electron transfer flavoprotein (ETF) complex to transfer electrons to complex III in the electron transport chain ^30^ (Fig. 4e). *L. monocytogenes* and *C. rodentium* infection decreased the abundance of the mRNAs of the ETF complex (Fig4. h), as well as transcripts encoding complex III (Fig. S5c), potentially further impacting GCDH enzyme activity. In addition, the level of glutarate is controlled by SUGCT, a glutarate-CoA transferase that converts glutarate to its CoA form to prevent excretion mediated carbon loss ^31^. Both infections decreased the transcript levels of SUGCT (Fig. 4i), supporting the idea that reduced abundance of GCDH and SUGCT may explain elevations of glutarate in bile of infected animals (Fig.1i, j). Host pathways for the derivation of 2-methylsuccinate are not clear, but this dicarboxylate may in part be derived from the microbiota ^32^.

In contrast to the other dicarboxylates where reduction of key transcripts appears to account for their induction by infection, *L. monocytogenes* and *C. rodentium* infection significantly stimulated the hepatic expression of *Acod1* (Fig. 4j), the gene encoding the itaconate-synthesizing enzyme aconitate decarboxylase 1 (Acod1, aka Irg1). Acod1 is mainly expressed in innate immune cells and converts the TCA cycle intermediate aconitate to itaconate (Fig. 4a). RNA-FISH based detection of *Acod1* transcripts in *C. rodentium* infected animals revealed markedly increased *Acod1* signal in the liver (Fig. 4k). Collectively, these data support the hypothesis that enteric infections shape the bile metabolome by modulating the expression of hepatic metabolic enzymes.

### Itaconate regulates the abundance of intestinal tuft cells

Since itaconate abundance was elevated in the bile of both *L. monocytogenes* and *C. rodentium* infected animals (Fig. 1i, j, Fig. S2i) and Acod deficient mice are available ^33^, we focused our analyses on unveiling the functions of this dicarboxylate in intestinal homeostasis and defense. While itaconate has known immunoregulatory activities outside of the intestine^34^, including suppression of inflammasome activation in bone-marrow derived macrophages ^35, 36^, the roles of itaconate in the intestine have not been explored. One of itaconate’s metabolic effects in immune cells is inhibition of the activity of the tricarboxylic acid cycle enzyme succinate dehydrogenase, leading to accumulation of succinate in culture supernatants ^37^. Concordant with these cell cultured-based observations, we found that the abundance of succinate in the bile of WT mice was significantly higher than in Acod1^-/-^ littermates following *C. rodentium* infection (Fig. 5a). Since succinate promotes the proliferation of intestinal tuft cells ^38^, we tested whether itaconate regulates the abundance of these chemosensory cells by comparing the number of tuft cells in Acod1^+/+^ and Acod1^-/-^ littermates. Acod1^+/+^ mice had ∼5-fold greater tuft cell abundance than Acod1^-/-^ mice in their ilea (Fig. 5b-c). Together, these observations are consistent with a model that itaconate stimulation of biliary and/or intestinal succinate levels elevates the abundance of intestinal tuft cells, potentially by signaling through the intestinal epithelial cell surface G-protein coupled receptor Sucnr1, which binds succinate ^39^.

**Figure 5.**
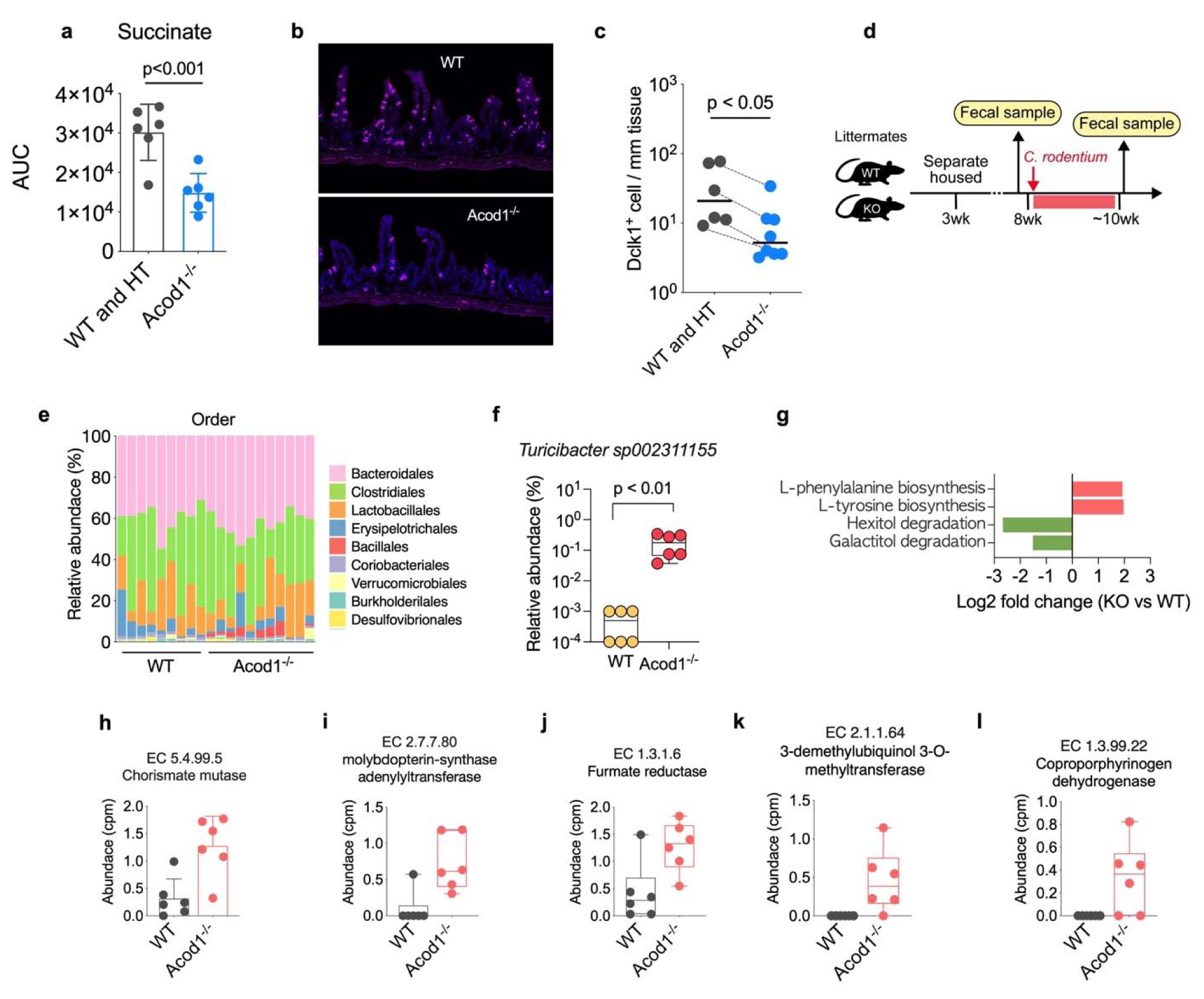
Itaconate regulates tuft cell abundance, and microbiota composition and functional potential respectively. (a) Abundance of succinate in bile of WT and Acod1^-/-^ mice post *C. rodentium* infection. (n=6 for WT and n=6 for Acod1^-/-^ mice) (b) Representative image of distal ileum of littermates of Acod1^+/+^ and Acod1^-/-^ mice treated with streptomycin, which elevates ileal succinate levels. Tuft cells were identified with anti-Dclk1 antibody (red) and nuclei were stained with DAPI (blue). (c) Quantification of tuft cell number in distal ileum. (Data from 4 litters of mice in 3 independent experiments, n=6 for WT, and n=8 for Acod1^-/-^ mice; lines connect littermates). (d) Experimental scheme for studying the effects of *Acod1* on fecal microbiota composition. Littermates of Acod1 mice from heterozygotes parents were separate-housed according to genotype at 3-weeks of age and challenged with *C. rodentium* at 8-weeks of age. Fecal samples were collected before and after clearance of *C. rodentium* infection; (the course of *C. rodentium* infection is ∼12-14 days). (e) Relative abundance of fecal microbiota at the order level post clearance of *C. rodentium* infection. (f) Abundance of *Turicibacter sp002311155* in WT and Acod1^-/-^ mice post *C. rodentium* infection identified by shotgun metagenomics (n=6 for WT, and n=8 for Acod1^-/-^ mice). (g) Differentially abundant pathways in Acod1^-/-^ mice compared to WT mice post *C. rodentium* infection. (h-l) The five most-enriched Enzyme Commission (EC) families in Acod1^-/-^ mice compared to WT mice post *C. rodentium* infection.

### Itaconate regulates the composition and function of the gut microbiota

Several bile components, including bile acids regulate both the abundance and function of intestinal commensal species^40, 41^. We investigated if itaconate regulates gut microbiota composition. In these experiments, Acod1^+/+^ and Acod1^-/-^ littermates were separately housed according to genotype when they were 3 weeks old and the fecal microbiota composition in the two groups was compared when the animals were ∼8 weeks old (Fig. 5d). 16S rRNA gene profiling showed that several OTUs, including two *Ruminococcaceae* species and one *Clostridiales* species showed marked difference in abundance in the Acod1^-/-^ and Acod1^+/+^ mice (Fig. S6a-d), suggesting that itaconate influences gut microbiota composition at steady state.

Since enteric infections disrupt the intestinal microbiota ^21^ and we found that the abundance of itaconate in bile increased with enteric infection (Fig. 1i, j), we investigated if itaconate impacts the recovery of the gut microbiota following *C. rodentium* infection. Acod1^-/-^ and Acod1^+/+^ mice were challenged with *C. rodentium* and we profiled their fecal microbiota during the recovery phase following pathogen clearance (Fig. 5d). The Acod1^-/-^ animals had a marked increase in the abundance of the *Bacillales* order post *C. rodentium* infection (Fig. 5e) that was mostly due to a >100-fold increase in the abundance of two *Bacillaceae* species (Fig. S6e, f, g). Shotgun metagenomics enabled identification of one of the species as *Turicibacter sp002311155*, whose abundance was ∼300-fold higher in Acod1^-/-^ mice (Fig. 5f). In contrast, the abundance of one *Clostridiales* species was significantly lower in Acod1^-/-^ mice (Fig. S6e, h), suggesting that itaconate helps to maintain its abundance during and/or following perturbations such as *C. rodentium* infection.

We leveraged the shotgun metagenomics data to understand how itaconate impacts the functional output of the gut microbiome following *C. rodentium* infection. Comparing to WT mice, 27 differentially abundant functions (Enzyme commission, EC) were identified in Acod1^-/-^ mice (Fig. S6i). Among them, chorismate mutase (EC 5.4.99.5), a key enzyme for both tyrosine and phenylalanine biosynthesis was enriched in Acod1^-/-^ mice (Fig. 5h); pathway abundance analysis also unveiled the elevation of these aromatic amino acid biosynthesis pathways in Acod1^-/-^ mice (Fig. 5g). Additionally, the abundance of functions that are critical for heme synthesis (Fig. 5l), and microbial anaerobic and aerobic respiration, including enzymes involved in ubiquinone synthesis (Fig. 5k), molybdopterin-synthase activation (Fig. 5i), and fumarate reduction (Fig. 5j) were elevated in Acod1^-/-^ mice, suggesting that itaconate inhibits specific functional pathways. Collectively, these data support the idea that itaconate modulates the composition and functional output of the gut microbiome following perturbations such as enteric infection.

### Itaconate promotes host defense against *Vibrio cholerae*

Itaconate has antimicrobial activity against several bacterial pathogens, including *Mycobacterium tuberculosis* and *Salmonella typhimurium*. ^42, 43^ To assess the role of itaconate in host defense against an extracellular enteric pathogen, we first challenged Acod1^-/-^ and Acod1^+/-^ littermates with *C. rodentium* and found no difference in fecal shedding of this pathogen (Fig. S7a). Since bile metabolites are likely to have higher concentrations in the small intestine than in the colon, we challenged neonatal Acod1^-/-^ and Acod1^+/-^ littermates with a pathogen that primarily replicates in the small intestine, *Vibrio cholerae*, the agent of cholera. ^44^ Comparing to Acod1^+/-^ mice, there was an ∼5 times greater *V. cholerae* burden recovered from the both proximal and distal small intestine of Acod1^-/-^ mice (Fig. 6a), suggesting that itaconate restricts *V. cholerae* growth within the small bowel.

**Figure 6.**
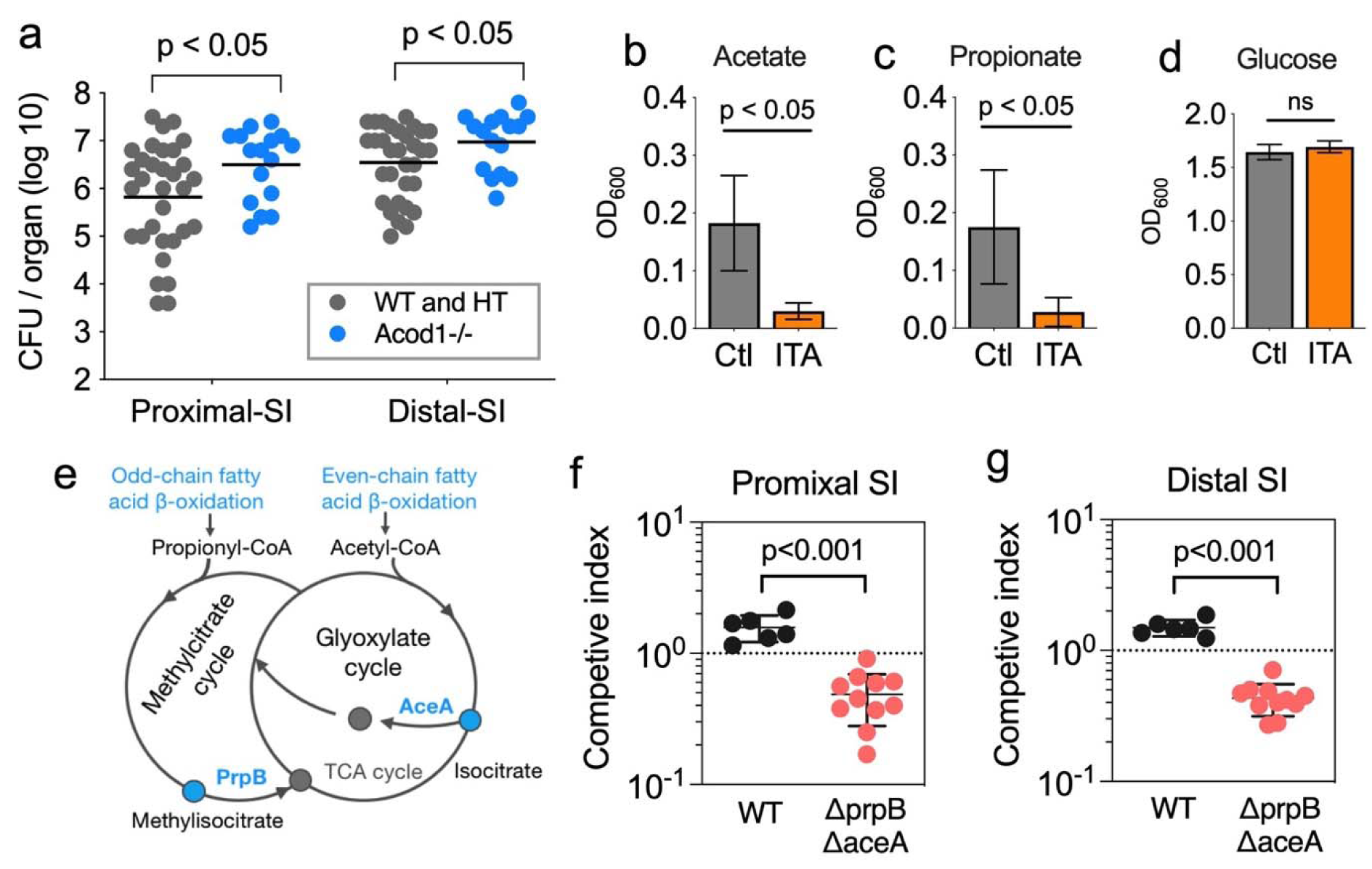
Itaconate confers resistance to *V. cholerae* intestinal colonization. (a) CFU of *V. cholerae* in the proximal and distal small intestine of infected WT and Acod1^-/-^ mice (Data from 4 independent experiments, n=32 for WT, and n=16 for Acod1^-/-^ mice) (b-d) Growth of *V. cholerae* in M9 minimal medium supplemented with 10mM acetate (b), propionate (c), or glucose (d) as the sole carbon source. (e) Scheme for acetate and propionate utilization in *V. cholerae*; AceA (isocitrate lyase) PrpB (methylisocitrate lyase). *aceA* vs the WT strain in proximal (f) and distal Δ (g) small intestine of infant mice.

The impact of itaconate on *V. cholerae* growth in a variety of *in vitro* conditions was assessed to test whether this dicarboxylate could directly inhibit the pathogen’s growth. Itaconate inhibits *M. tuberculosis* growth on acetate and propionate by impairing the activity of the pathogen’s isocitrate lyase and methyl-isocitrate lyase, ^45^ enzymes that mediate utilization and/or detoxification of short chain fatty acids. Itaconate inhibited *V. cholerae* growth when acetate or propionate were used as the sole carbon source (Fig. 6b, c), but not when glucose was the carbon source (Fig 6d). These data support the idea that itaconate inhibits the activities of *V. cholerae* isocitrate lyase (AceA) and methyl-isocitrate lyase (PrpB), curbing the glyoxylate (GC) and methyl-isocitrate (MC) cycles that govern utilization of even and odd chain fatty acids, respectively (Fig. 6e).

Since fatty acid utilization is thought to support robust *V. cholerae* intestinal colonization of infant mice, ^46^ we tested whether AceA and PrpB contributed to *V. cholerae* growth in the infant mouse intestine. A *V. cholerae* mutant strain (Δ*aceAΔprpB*) lacking both isocitrate lyase and methyl-isocitrate lyase exhibited a competitive colonization defect (Fig. 6f, g), supporting the idea that the pathogen relies on fatty acids for robust growth in the host intestine. Collectively, these data are consistent with the hypothesis that itaconate mediates host defense against *V. cholerae* by impairing the pathogen’s capacity to consume fatty acids in the intestine.

### 2-methylsuccinate suppresses intestinal inflammasome activity

Itaconate suppresses inflammasome activation in bone-marrow derived macrophages ^35, 36^. We tested whether addition of this dicarboxylate to the drinking water of uninfected SPF mice also regulated inflammasome activity in the proximal small intestine. However, itaconate supplementation did not change IL-18 production in an *ex vivo* system (Fig. S7c). Recently dicarboxylates, including itaconate have been found to be capable of supporting the respiration of gut microbiota ^32^. In *Eggerthella lenta*, respiration of itaconate yields 2-methylsuccinate ^32^, but little is known about the function of this dicarboxylate, which was elevated in the bile of infected animals (Fig. 1i, j). Mice that received 2-methylsuccinate in their drinking water had significantly less IL-18 released into supernatants from duodenal explants compared to control animals (Fig. S7c), suggesting that 2-methylsuccinate suppresses intestinal inflammasome activity and raising the possibility that additional bile metabolites have immunomodulatory activity. Collective, itaconate and its downstream products, including 2-methylsuccinate, appear to control the composition of the intestinal epithelium and microbiota, as well as intestinal immune function and defense.

## Discussion

Bile has been recognized as a vital body fluid important for the maintenance of health at least since the days of Hippocrates ^47^.. Here, we profiled the murine bile metabolome using untargeted metabolomics to create a comprehensive atlas of bile metabolites and investigated how the microbiota and enteric infection modulate the bile metabolome to gain insights into bile function. Our findings provide a new perspective on bile function. We discovered that the bile metabolome is not only more complex than was appreciated but is shaped by the microbiota and re-modeled by enteric infection. Moreover, we found that an infection-stimulated bile metabolite modulates the composition of the intestinal epithelium and microbiota as well as intestinal immune function and defense. Changes in hepatic transcriptional profiles induced by enteric infection likely explain how infection re-models bile composition. Together, our findings unveil a new interorgan defense circuit embedded in entero-hepatic circulation in which intestinal infection stimulates modifications in bile composition that in turn modulate intestinal function (Fig. S8).

Our observations suggest that enteric infection leads to release of signals that enable the intestine to leverage the capacity of liver to synthesize bile in order to modulate gut functions including immune regulation and defense (Fig. S8). The delivery of infection-stimulated modified bile to the intestine through the common bile duct can be thought of as analogous to the acute phase response, where infection-stimulated hepatic products are secreted into the blood to aide in systemic defense and tissue repair ^4^. However, modification of bile composition in response to enteric infection targets the intestine and represents a novel innate defense circuit to guard the intestine that functions along with autonomous enteric defense systems ^48^. It is possible that systemic infection also triggers changes in bile composition and that the liver’s acute phase response routinely has biliary as well as systemic outputs.

The liver responds to enteric infection even in the absence of detectable pathogen cells replicating in liver (Fig. 3a and S3a), suggesting that the hepatic sensing mechanisms are at least in part driven by host-derived factors, such as cytokines, along with microbial-associated products, such as LPS or peptidoglycan. However, although no recoverable *C. rodentium* was found in some animals with changes in hepatic transcriptional responses at ten days post-infection, it is possible that pathogen cells were cleared from the liver by that point. Infection-associated signals are presumably delivered to the liver primarily via the portal circulation, but some role for the systemic circulation is also possible. The liver is known to monitor the abundances of bile acids in the systemic circulation in order to modulate the synthesis of primary bile acids ^12^; hepatic sensing of infection-induced signals to trigger modifications in bile composition may be an analogous process. Identifying the signals and hepatic sensing and regulatory elements that govern the type, magnitude and duration of hepatic driven changes in bile makeup are key challenges for future studies.

*C. rodentium* and *L. monocytogenes* infection led to rewiring of hepatic metabolic circuitry and these changes likely underpin many changes in bile composition. Remodeling of the liver’s metabolic output, particularly in arachidonic acid and cholesterol metabolism was also observed in animals infected with *Salmonella typhimurium* ^49^. Infection-related signals, such as cytokines, presumably activate metabolic switches, such as Hk2, that mediate the shift from aerobic respiration to fermentation, as we observed in enteric infections. Such switches alter cell function in order to enhance cellular adaptation to perturbation (e.g., increased effector production) ^50^, promoting organismal recovery and survival. Hepatic detoxification systems appeared to largely shut down during enteric infection, as has been observed in LPS and TNF ^51, 52^. The physiologic ramifications of the reduction in detoxification warrant further investigation; however, it seems likely that this response favors allocation of resources to defense programs that directly support host survival.

Most studies of dicarboxylates have addressed issues related to their intracellular activities. The four dicarboxylates whose abundance in cell-free bile increased in both *C. rodentium* and *L. monocytogenes* infection were not known to be bile constituents. Thus, our findings suggest that 2-methylsuccinate, which suppressed intestinal inflammasome activity, and itaconate, which modified the composition of the intestinal epithelium and the gut microbiota and curbed *V. cholerae* intestinal colonization, can function extracellularly. Regulatory functions of itaconate and glutarate have been attributed to their ability to covalently modify mitochondrial and cytosolic proteins ^53, 54^, and if biliary itaconate has a similar mechanism, then there must be pathways for its uptake into intestinal cells. Import of itaconate into cytotoxic CD8 T cells has been demonstrated ^55^, but specific mechanisms facilitating cell entry of this TCA cycle metabolite across cell membranes remain to be described. Import of additional dicarboxylates may underlie their regulatory effects on host and potentially microbial cells as well. Alternatively, the metabolites may function extracellularly by binding to target cell surface receptors, similar to the binding of succinate and indole-3-acetic acid to G-protein coupled receptors ^39, 56^. Recently dicarboxylates, including itaconate and fumarate have been found to be capable of supporting the respiration of gut microbiota and the effects of these dicarboxylates on microbial physiology warrant further investigation.

Itaconate is known to have immunomodulatory activity and is primarily thought to impede inflammation. We found that 2-methylsuccinate also has inflammasome-suppressive activity. We speculate that 2-methylsuccinate, and perhaps itaconate as well, control the magnitude and duration of gut inflammation following enteric infection, and thus represent host ‘tolerance’ factors that promote the maintenance of intestinal tissue integrity during infection. Investigations of the functions of glutarate and 2-oxoadipate in modulating intestinal homeostasis will require additional knowledge of the genes and processes that govern their abundance.

Our findings also extend knowledge of the intricate connection between pathogen metabolic preference and infection outcome ^57^. We found that itaconate promotes host defense against *V. cholerae*, linking this metabolite to host defense against an extracellular pathogen in the intestinal tract. Acod1-/- mice are not only deficient in bile-derived itaconate, and other sources of this dicarboxylate (e.g. neutrophils) could contribute to the elevated *V. cholerae* growth in Acod1-/- mice; however, innate immune cells are not thought to play a major role in defense against this pathogen ^58^. Acod1-/- mice have heightened susceptibility to *M. tuberculosis but not to Listeria monocytogenes* ^59^, indicating that itaconate mediated defense is not effective against all pathogens. The differential potency of itaconate vs different pathogens may be explained by differences in metabolic strategies that pathogens rely on *in vivo*. Itaconate inhibits key microbial enzymes (methylcitrate lyase and isomethylcitrate lyase) that facilitate pathogen fatty acid utilization. Both *M. tuberculosis* and *V. cholerae* rely on fatty acid metabolism to fuel in vivo growth ^46, 57^ likely explaining how itaconate restricts the growth of these pathogen. Given the rich fatty acid content in bile, the presence of itaconate in bile may be a host strategy to guard against pathogen utilization of this source of energy.

In sum, our findings uncovered the complex and dynamic nature of bile composition and expands knowledge of the homeostatic functions of the liver and bile, one of its major products. These observations underscore the profound influence that the liver exerts on intestinal tract functions. Mobilization of this entero-hepato-biliary regulatory circuitry may have translational applications in the treatment of a variety of intestinal disorders.

## Methods

### Animals

All animal experiments were conducted following the protocol (2016N000416) reviewed and approved by the Brigham and Women’s Hospital Institutional Animal Care and Use Committee. Special pathogen free (SPF) C57Bl/6J mice were purchased from the Jackson Laboratory (stock no. 000664). Mice were kept in Harvard Medical School animal facility for at least 72 hours prior to the experiments. Germ free (GF) C57Bl/6J mice were purchased from the Massachusetts Host-Microbiome Center. Acod1^-/-^ mice were purchased from the Jackson Laboratory (C57BL/6NJ-Acod1em1(IMPC)J/J, stock no. 029340) and bred in Harvard Medical School animal facility. C57Bl/6 with 3-day postnatal infants (P3) were purchased from the Charles River Laboratories (stock no. 027) and kept in Harvard Medical School animal facility until postnatal day-5 (P5). All mice were kept under the 12-hour light-dark cycles: lights being turned off at 7 p.m. and turned on at 7 a.m.

### Global metabolomic profiling of bile

Female SPF mice (9 to 10-week-old age) were orally inoculated with *Listeria monocytogenes* or *Citrobacter rodentium* following the methods described previously ^18, 61^. Briefly, mice were deprived of food for 6 hours, lightly sedated with isoflurane inhalation, and oro-gastrically inoculated with 3 ×10^9^ CFU of *Listeria monocytogenes* 10403S InlA^m^ strain in a 300 µl mixture of 200 mM CaCO3 in PBS, or with 1 ×10^9^ CFU *Citrobacter rodentium* ICC168 strain in 200 µl PBS using 18G flexible feeding needles (DT 9928, Braintree Scientific). Uninfected SPF mice were inoculated with 200 µl of PBS. Animals were sacrificed at 4 days post *L. monocytogenes* and 10 days *C. rodentium* infection, respectively. Bile samples were collected from the gallbladder using insulin syringes (BEC-309311, Becton Dickinson), filtered with 0.22 µm centrifuge tube filters (8160, Corning), snap frozen in liquid nitrogen, and stored in −80 °C until analysis. To obtain a minimal volume of 60 µl of bile for global metabolomic profiling, samples were generated by pooling bile from 3-5 SPF mice, 3-4 *L. monocytogenes* or *C. rodentium* infected mice, and 2-3 GF mice. The liver and bile of *L. monocytogenes* infected mice and the liver and colon of *C. rodentium* infected animals were used for bacterial burden enumeration.

Bile samples were processed and analyzed by Metabolon (Metabolon Inc., Morrisville, NC) for global metabolomic profiling. To remove protein, dissociate small molecules bound to protein or trapped in the precipitated protein matrix, and to recover chemically diverse metabolites, proteins were precipitated with methanol under vigorous shaking for 2 min (Glen Mills GenoGrinder 2000) followed by centrifugation. The resulting extract was divided into four fractions for analysis: two for analysis by two separate reverse phase (RP)/UPLC-MS/MS methods with positive ion mode electrospray ionization (ESI), one for analysis by RP/UPLC-MS/MS with negative ion mode ESI, one for analysis by HILIC/UPLC-MS/MS with negative ion mode ESI. Raw data was extracted, peak-identified and QC processed using Metabolon’s hardware and software. Peaks were quantified using area-under-the-curve and reported as “Original Scale” data, and significance were determined by Welch’s two-sample *t*-test. For metabolite clustering, “Original Scale” data was used for calculating Z-score. The clustering of the metabolites was performed using pam (Partitioning Around Medoids) function in the R-package cluster (v2.1.4) with a parameter ‘k=7’.

### Quantification of bile dicarboxylates

For quantification of abundance of itaconate, 2-methylsuccinate, glutarate, and 2-oxoadipate (i.e., four dicarboxylates) in bile, female SPF mice (9 to 10-week-old age) were oro-gastrically inoculated with *C. rodentium* or PBS as described above. Bile samples were collected from individual infected mouse, processed as described above, and prepared for LC-MS/MS analysis following the sample preparation guideline ^62^. Briefly, one volume (8 µl) of bile samples was combined with four volume of cold methanol (-80 °C), gently mixed, and incubated at −80 °C for 6 hours. Samples were centrifuged at 14,000 g for 10 min (4 °C), and supernatants were transferred to new tubes and lyophilized to pellets using no heat. Abundance of four dicarboxylates in samples were quantified using 5500 QTRAP LC-MS/MS system at the mass spectrometry core facility at Beth Israel Deaconess Medical Center. Commercially available glutarate (G3407, Sigma), itaconate (I29204, Sigma), 2-methylsuccinate (AAH6096714, Fisher), and 2-oxoadipate (75447, Sigma) were used as chemical standards. Peaks were quantified using area-under-the-curve and statistical significance was determined by Mann-Whitney test.

### RNA-seq analysis of liver

C57BL/6J mice were infected with *L. monocytogenes* or *C. rodentium* following the protocols described above. RNAs were extracted from the liver samples using Trizol (Thermo Fisher) and RNA-seq libraries were prepared by KAPA RNA Hyperprep Kit (Roche). The libraries were sequenced on an Illumina NextSeq 550 instrument with paired-end runs of 2×75 bp. The reads were mapped to the mouse reference genome (mm10) using STAR v2.7.3a with default parameters. Differentially expressed genes were identified using the R-package DESeq2 (v1.36.0) with adjusted p-value < 0.05, and the clustering was performed using pam (Partitioning Around Medoids) function in the R-package cluster (v2.1.4) with a parameter ‘k=7’. Pathway analysis was performed by the R-package clusterProfiler (v4.4.4) with gene sets from msigdbr R-package (v7.5.1) or KEGG metabolism pathways.

### Metabolic network analysis (integrated analysis of RNA-seq and metabolites)

Metabolic network analysis was performed using Shiny GATOM (https://artyomovlab.wustl.edu/shiny/gatom/) with parameters ‘Network type=KEGG network’, ‘Network topology = atoms’, ‘Scoring parameter for genes=P-value threshold’, ‘Scoring parameter for metabolites = P-value threshold’ and the thresholds = -4. The fold changes in input data for genes were shunk using the function lfcShrink (type=”apeglm”) in the R-package DESeq2. The fold changes for metabolites were “Fold of Change” of *L. monocytogenes*-or *C. rodentium*-infected mice compared to SPF mice reported by Metabolon.

### Measurement of IL-18 production in duodenal explants

Mice were provided with 5 mM of itaconate (I29204, Sigma) and 2-methylsuccinate (AAH6096714, Fisher) in their drinking water individually (adjusted to pH 7 using NaOH) for 8 days. Sodium concentration matched water (NaCl, S5150, Sigma) was used as control. After euthanasia, 3-cm of the proximal intestine tissues were collected, opened longitudinally, weighted, and transferred to 1 ml of DMEM with glucose and glutamine (11965092, Gibco), 10% heat-inactivated FBS (Gibco), and penicillin and streptomycin (15140122, Gibco) as previously described ^60^. The intestinal explants were incubated in a 37°C cell culture incubator with 5% CO_2_ for 20 hours. Explant culture media was centrifugated at 12,000 g for 5 min, and supernatants were collected and stored at −80 °C. Abundance of IL18 were measured using ELISA kit (7625, R&D systems) and normalized per milligram of tissue.

### Quantification of bile succinate

Littermates of adult female mice (10 to 14-week-old) from Acod1^+/-^ parents were infected with *C. rodentium* following the protocol described above and euthanized at 10 day-post infection. Bile samples were collected from individual infected mouse and 4 µl of bile were used for sample preparation following the protocol described above (quantification of bile dicarboxylates). Abundance of succinate in samples were quantified using a polar metabolite detection pipeline and 5500 QTRAP LC-MS/MS system at the mass spectrometry core facility at Beth Israel Deaconess Medical Center.

### Tuft cell immunohistology

Littermates of Acod1^-/-^, Acod1^+/-^, and Acod1^+/+^ adult male mice (10 to 16-week-old) were used for experiments. Mice were administered with 200 µl of streptomycin (20mg) in water daily via oral gavage using 18G flexible feeding needles (DT 9928, Braintree Scientific) for 5 days and euthanized at day 7 ^38^. 1.5-cm-long distal ileum tissues were dissected, fixed with 4% PFA for 2h at room temperature, and transferred to 30% sucrose in PBS at 4 °C overnight. The samples were embedded in 1:2.5 of 30% sucrose and OCT solution and cut into 10 µm sections. The slides were washed three times with PBS and block with 5% normal goat serum with 0.3% Triton X-100 for 1 h at room temperature. Slides were incubated with Rabbit anti-DCAMK polyclonal antibody (1:500 dilution, ab31704, Abcam) in 4 °C overnight, and Alexa-594 Goat anti-Rabbit IgG secondary antibody (1:1000 dilution, A-11072, Thermo Fisher) for 1 hour at room temperature. DNA was labeled with DAPI (P36935, Thermo Fisher). Images were captured using Leica Stellaris Confocal Microscope at the Microscopy Resources On the North Quad (MicRoN) core of Harvard Medical School. For quantification of tuft cell frequency, the number of Dclk1 positive cells were enumerated in a 2.5 mm-long representative tissue and presented as number of tuft cells per mm tissue.

### Analysis of fecal microbiota composition

Littermates of female mice from Acod1^+/-^ parents were separate-housed according to their genotypes (Acod1^-/-^ vs Acod1^+/-^ and Acod1^+/+^) at 3-week-old age and infected with *C. rodentium* at 8-week-old age following the protocol described above. Fecal pellets were collected at 1 day prior to *C. rodentium* infection and post pathogen-clearance (∼14 days post infection). The genomic DNA were extracted from fecal samples following the method described previously ^63^. V3-V4 region of 16S rRNA was amplified using primer 341F and 805R and libraries were prepared using Nextera XT Index Kit v2 (Illumina). The libraries were sequenced on Illumina Mi-Seq instrument using Miseq Reagent Kit v3 (600 cycles) with paired-end runs. Qiime2 pipeline ^64^ were used to process the reads and Greengene reference library was used for taxonomy mapping. Differential abundance of OTUs were analyzed using R package ANCOMBC ^65^.

For shotgun metagenomic sequencing, DNA samples prepared as described above and sent for sequencing in SeqCenter (Pittsburgh, PA) using 2×150 cycles paired-end runs. Data were processed for quality control using KneadData ^66^. For taxonomical profiling, reads were processed using a k-mer method by Kraken2 ^67^ using a Kraken 2 database from The Mouse Gastrointestinal Bacterial Catalogue (MGBC) project ^68^. Taxonomical abundance was estimated using Bracken ^69^. Microbial functional profiling was performed using Humann3 ^66^.

### *In vitro* growth inhibition of *Vibrio cholerae* by itaconate

Growth of *V. cholerae* in defined nutrient conditions were performed using M9 minimal medium containing 1 mM MgSO4, 0.3 mM CaCl2, and one carbon source as indicated: acetate (10 mM) or glucose (0.5%). 1x trace elements solution was added when propionate (10 mM) was used as carbon source. To test the growth inhibitory effect of itaconate, culture mediums were supplemented with a final concentration of 10 mM itaconate and adjusted pH to 7 using NaOH. *V. cholerae* colonies from a LB plate were washed twice with M9 minimal medium, adjusted to OD of 0.5, and 5 µl of prepared *V. cholerae* culture were inoculated into 1ml medium of indicated conditions. All cultures were grown at 37 °C with shaking. OD were measured at 24 hours (for growth in acetate or glucose) or 96 hours (for growth in propionate) post inoculation.

### Infection of Acod1^-/-^ mice with *V. cholerae*

Littermates of infant mice at postnatal day 5 (P5) from Acod1^+/-^ breeding parents were orally inoculated with Haiti WT *V. cholerae* (total 10^5^ CFUs) in 50 µl LB, a dose that leads to robust intestinal colonization ^70^. Pups were euthanized at 20 hours post infection. Proximal and distal section of the small intestine were dissected, homogenized, and plated on LB Sm plates for CFU enumeration. Tail samples were used for genotyping.

### Construction of V. cholerae ΔaceAΔprpB variant

*V. cholerae* Haiti Δ*aceA*Δ*prpB* strain was created using allelic exchange method as previously described (ref). Briefly, the HaitiWT or Haiti Δ*aceA V. cholerae* strain was conjugated with MFDpir *E. coli* harboring the suicide plasmid pCVD442 that carries the upstream and downstream genomic region (∼700 bp each) of the targeted gene. The donor and recipient strain were mixed at a 1:1 ratio and incubated at 37 °C for 4 h. To obtain the single cross overs, conjugated reactions were streaked on LB plates with Sm/Cb. Double cross overs were isolated by restreaking single cross-over colonies on LB plates with 10% sucrose and incubating at room temperature for 2 days. Duplicate patching was used to examine the Cb resistance of the colonies and the correct double cross-over were identified by screening Sm R /Cb S colonies using colony PCR.

### V. *cholerae* competition assay

Overnight cultures of *V. cholerae* Haiti WT *lacZ+* and Δ*aceA*Δ*prpB lacZ-* strain were washed once with PBS and diluted 1:1000 in PBS. Pups at postnatal day 5 (P5) were orally inoculated with a 1:1 mixture (total 105 CFUs) of WT and Δ*aceA*Δ*prpB V. cholerae* strain in 50 μl PBS. Animals were euthanized at 20 hours post infection. Proximal and distal section of small intestine were dissected, homogenized, and plated on LB + Sm/X-gal for blue/white colony counting. Competition index (CI) was generated by dividing the ratio of white:blue Δ *prpB* /WT) colonies in the SI by the ratio of white:blue colonies in the inoculum, and compared to CI of Haiti WT *lacZ-* and Haiti WT *lacZ+* strain.

### RNAscope of *Acod1*

Liver samples from *C. rodentium* infected or PBS-treated animals were embedded in 1:2.5 of 30% sucrose and OCT solution and cut into 10 µm sections and stored at −80 °C. Staining following the fresh frozen tissue protocol provided by ACDbio. Briefly, the slides were fixed with 4% PFA that was pre-cold to 4 °C and incubated at 4 °C for 10 min and washed three times with PBS. Digested with protease IV and hybrid with Acod1 probe (stock 450241, ACDbio). Stain with Opal 570. Slides were incubated with Rabbit anti-CD19 polyclonal antibody (1:500 dilution, ab31704, Abcam) in 4 °C overnight, and Alexa-488 Goat anti-Rabbit IgG secondary antibody (1:1000 dilution, A-11072, Thermo Fisher) for 1 hour at room temperature. DNA was labeled with DAPI (P36935, Thermo Fisher). Images were captured using Leica Stellaris Confocal Microscope at the Microscopy Resources On the North Quad (MicRoN) core of Harvard Medical School.

## Acknowledgements

We acknowledge members of the Waldor laboratory and Dr. Brandon Sit for helpful discussions. We thank Rachel T Giorgio for assisting with RNAseq analysis, Dr. Bolutife Fakoya and Dr. Deborah R Leitner for assisting with infant mice experiments. Gnotobiotic mouse studies were supported by the Massachusetts Host-Microbiome Center with funding from the NIH (P30DK034854). Microscope analyses were supported by the Microscopy Resources On the North Quad (MicRoN) core of Harvard Medical School. This work was supported by NIH R01AI042347 grant to M.K.W., the Howard Hughes Medical Institute (HHMI) to M.K.W.

---

**Supplementary Figure 1.**
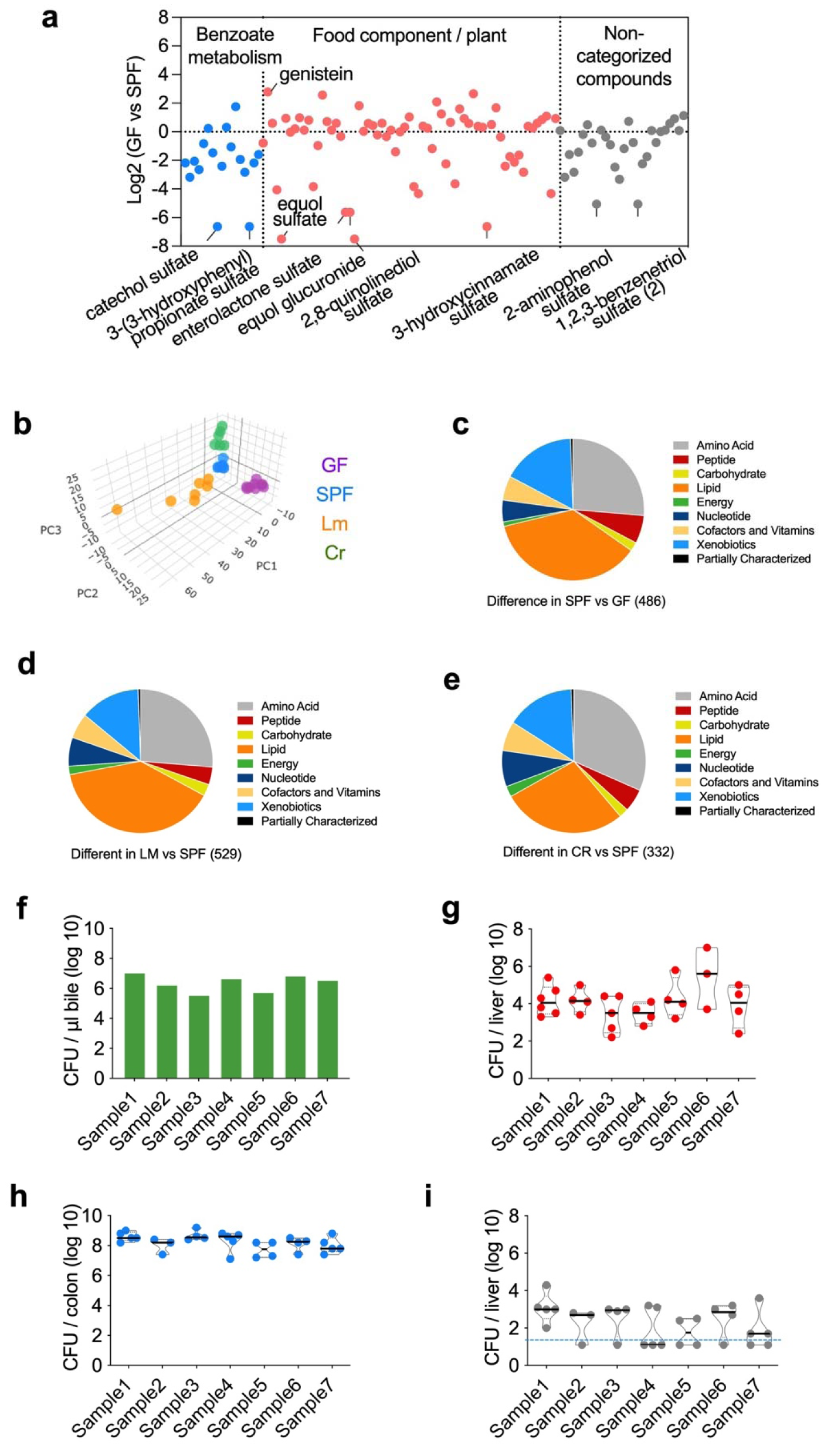
Microbiota and enteric infection alter the bile metabolome. (a) The relative abundance of metabolites in benzoate metabolism (blue), food components (pink), and non-categorized compounds category in bile of GF mice comparing to SPF mice. (b) Principal component analysis (PCA) of the bile metabolome in animals from the following four groups of mice: SPF, specific pathogen free, GF, germ free, Lm, *L. monocytogenes* infected, and Cr, *C. rodentium* infected. Each dot represents one sample. (c-e) The functional categories of metabolites with differential abundance in the indicated comparison. (f-i) The CFU of *L. monocytogenes* in bile (f) and liver (g) at 4-day post orogastric infection, and *C. rodentium* in colon (h) and liver (i) at 10-day post orogastric infection. The dotted line represents the limit of detection.

**Supplementary Figure 2.**
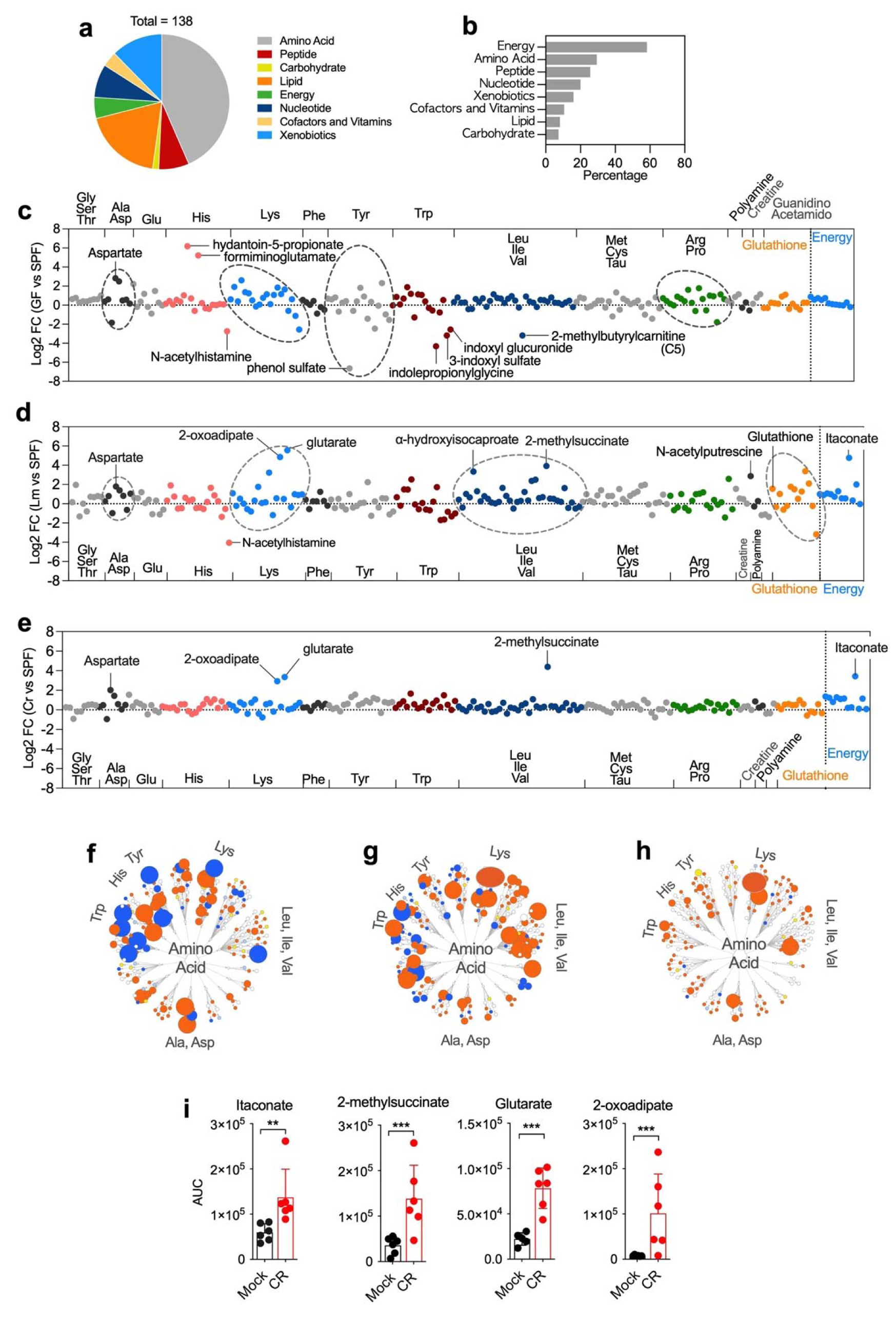
Microbiota and enteric infections alter the abundance of bile metabolites related to amino acids and energy. (a) The functional categories of bile metabolites that were elevated in both Lm and Cr infection. (b) The percentage of metabolites in each functional category that were elevated in both infection conditions. (c-e) The differential abundance of metabolites in the indicated categories. Comparisons of GF vs SPF (c), Lm vs SPF (d), and Cr vs SPF(e) are shown. (f-h) Changes in abundance of metabolites linked to amino acid metabolism pathways. GF vs SPF (f), Lm vs SPF (g), Cr vs SPF (h). Blue and orange represent decreased and increased abundance respectively; the size of the dots is related to the magnitude of the difference. (i) Quantification of the relative abundance of itaconate, 2-methylsuccinate, glutarate, and 2-oxoadipate in bile using LC_MS/MS.

**Supplementary Figure 3.**
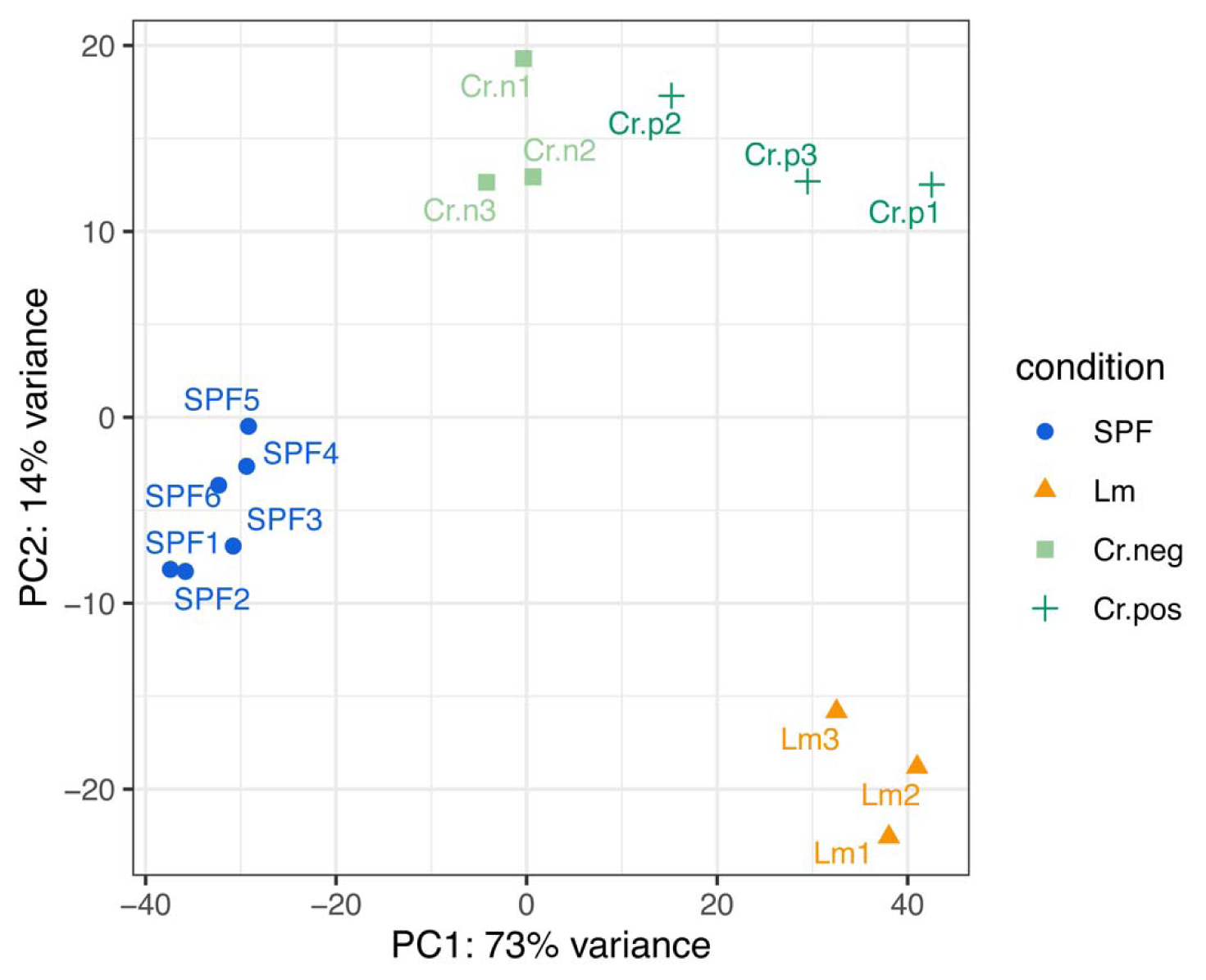
Enteric infections alter hepatic gene expression profiles. (a) Principal component analysis (PCA) analysis of liver transcriptomes in uninfected mice (SPF), Lm infected animals at 4 DPI, and Cr infected animals at 10 DPI. *C. rodentium* infected animals were divided into two groups, depending on whether or not they had detectable (Cr positive) or undetectable (Cr negative) *C. rodentium* cfu on the liver.

**Supplementary Figure 4.**
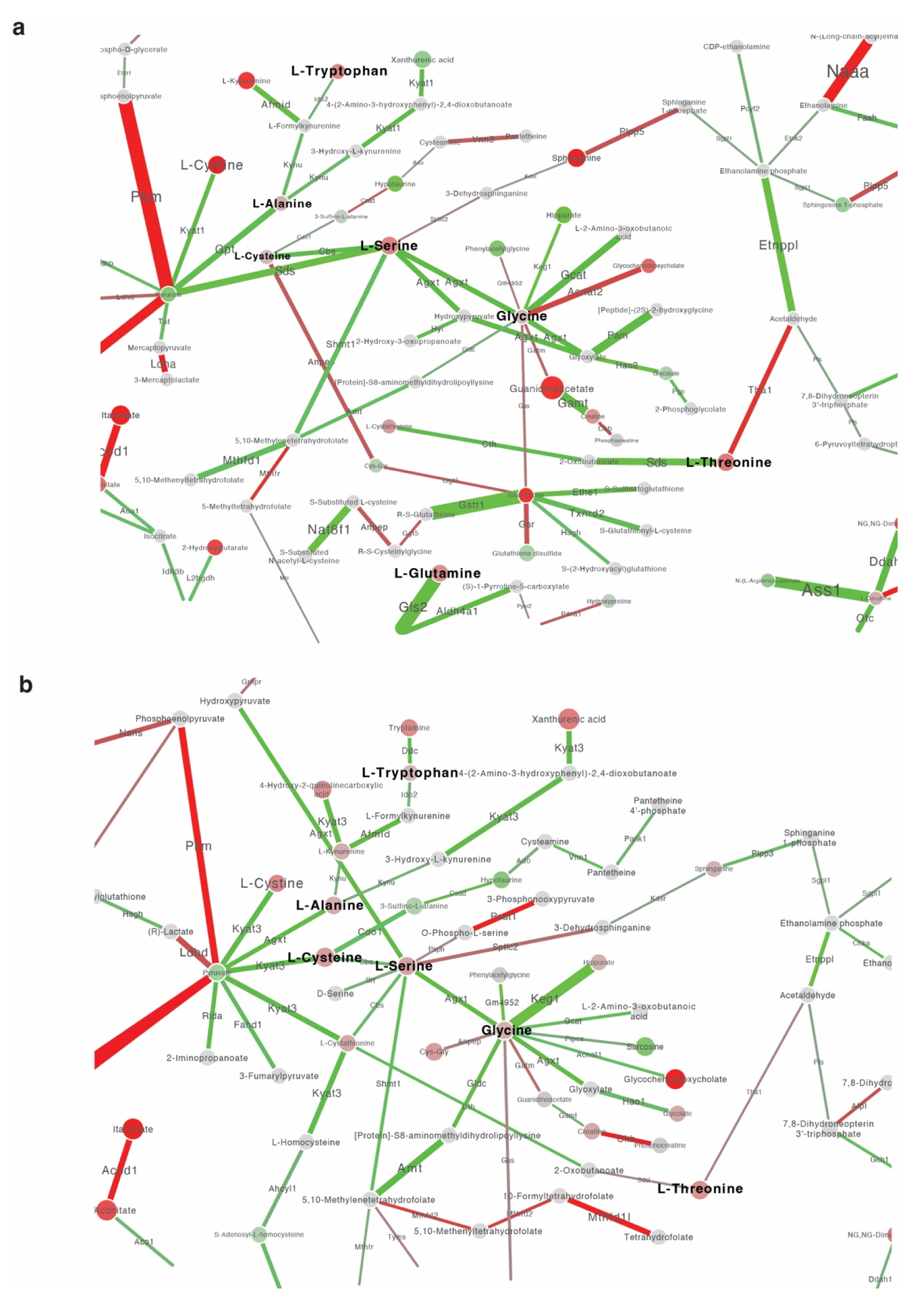
Network analysis of liver transcriptome and bile metabolome. Integrated analysis of genes and metabolites regulated by *L. monocytogenes* and *C. rodentium* infection was carried out using “Shiny GATOM: integrated analysis of genes and metabolites” (https://artyomovlab.wustl.edu/shiny/gatom/). The lines represent enzymes and the dots represent metabolites. The size of the dots and the thickness of the lines correlates with statistical significance and the color scale for the dots and lines represents fold change, where red color represents increased abundance and green color represents decreased abundance.

**Supplementary Figure 5.**
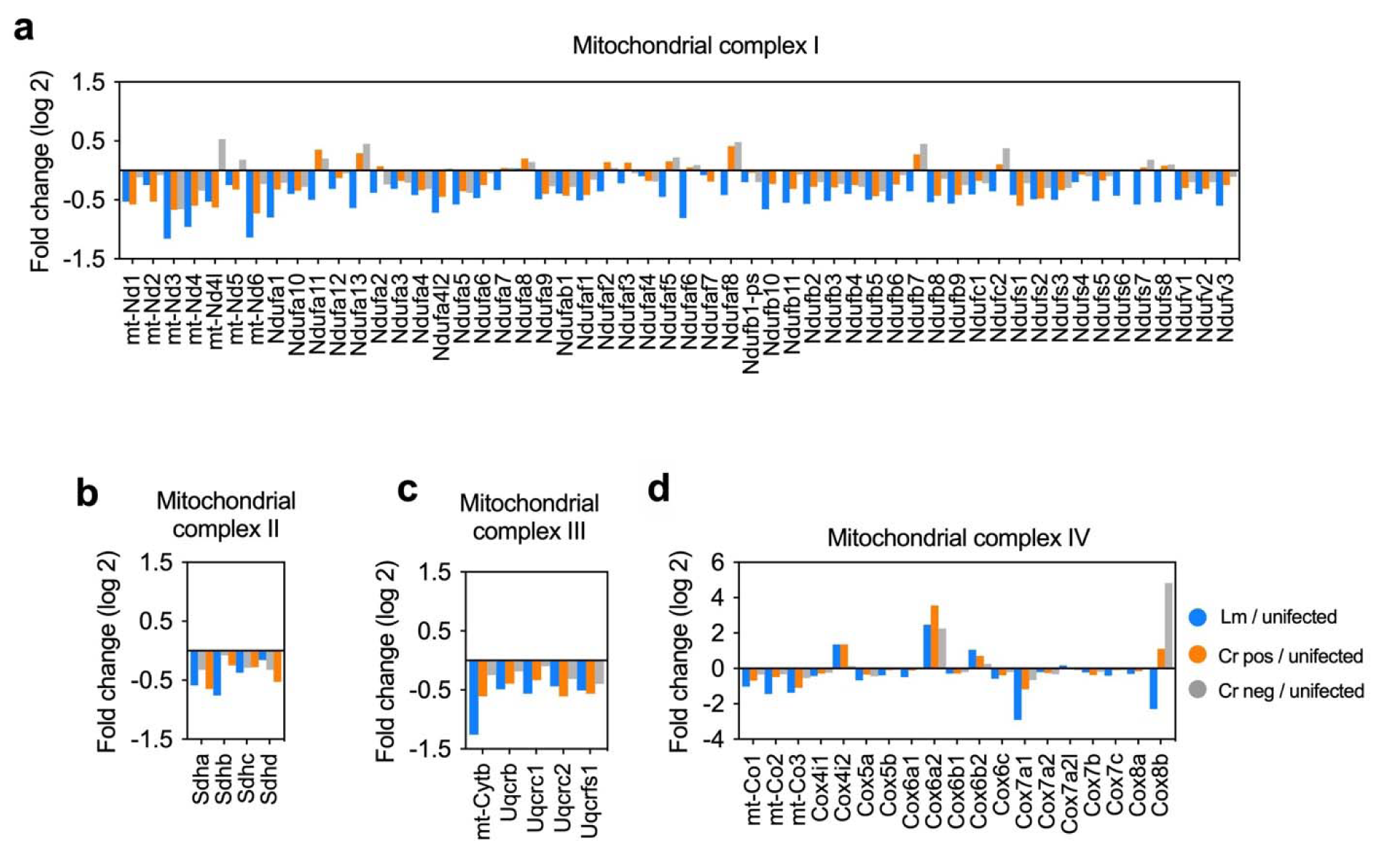
Transcripts for liver mitochondrial complexes I, II, III, and IV have reduced abundance during enteric infection. (a-d) Differential transcript abundance of liver mitochondrial complex I (a), complex II (b), complex III (c), and complex IV (d) in Lm infected, Cr infected liver positive or liver negative animals compared to SPF animals.

**Supplementary Figure 6.**
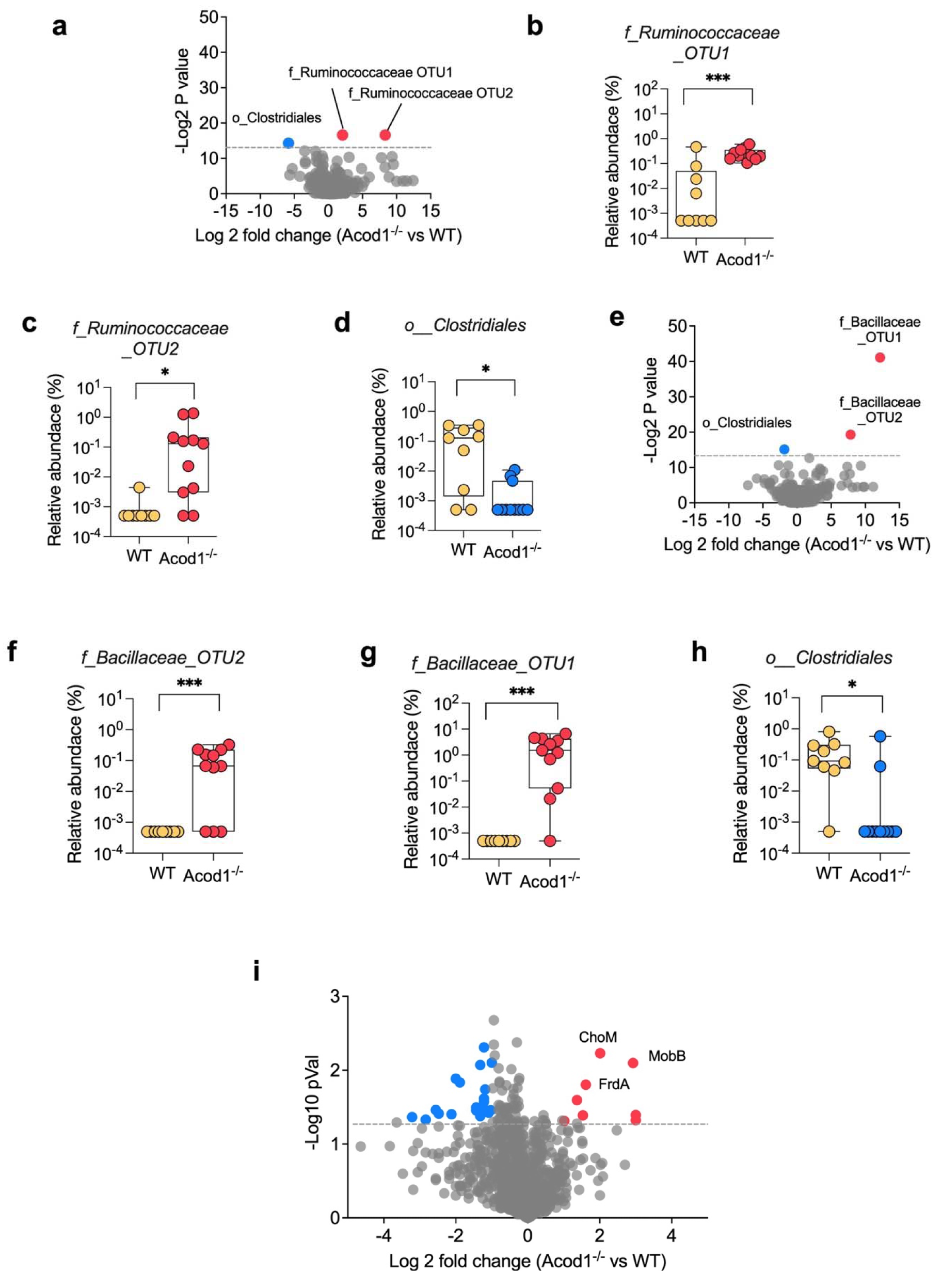
Enteric infection-stimulated bile metabolites regulate gut microbiota composition. a) Differential abundance of fecal microbiota at the OTU level between WT and Acod1^-/-^ mice before *C. rodentium* infection. (b-d) The three most differentially abundant OTUs in Acod1^-/-^ mice compared to WT mice before *C. rodentium* infection. (i) (e) Differential abundance of fecal microbiota at the OTU level between WT and Acod1^-/-^ mice post *C. rodentium* infection. (f-h) Three most differentially abundant OTUs in Acod1^-/-^ mice compared to WT mice post *C. rodentium* infection. (i) Differential abundance of functions (Enzyme Commissions, ECs) in WT vs Acod1^-/-^ mice post *C. rodentium* infection.

**Supplementary Figure 7.**
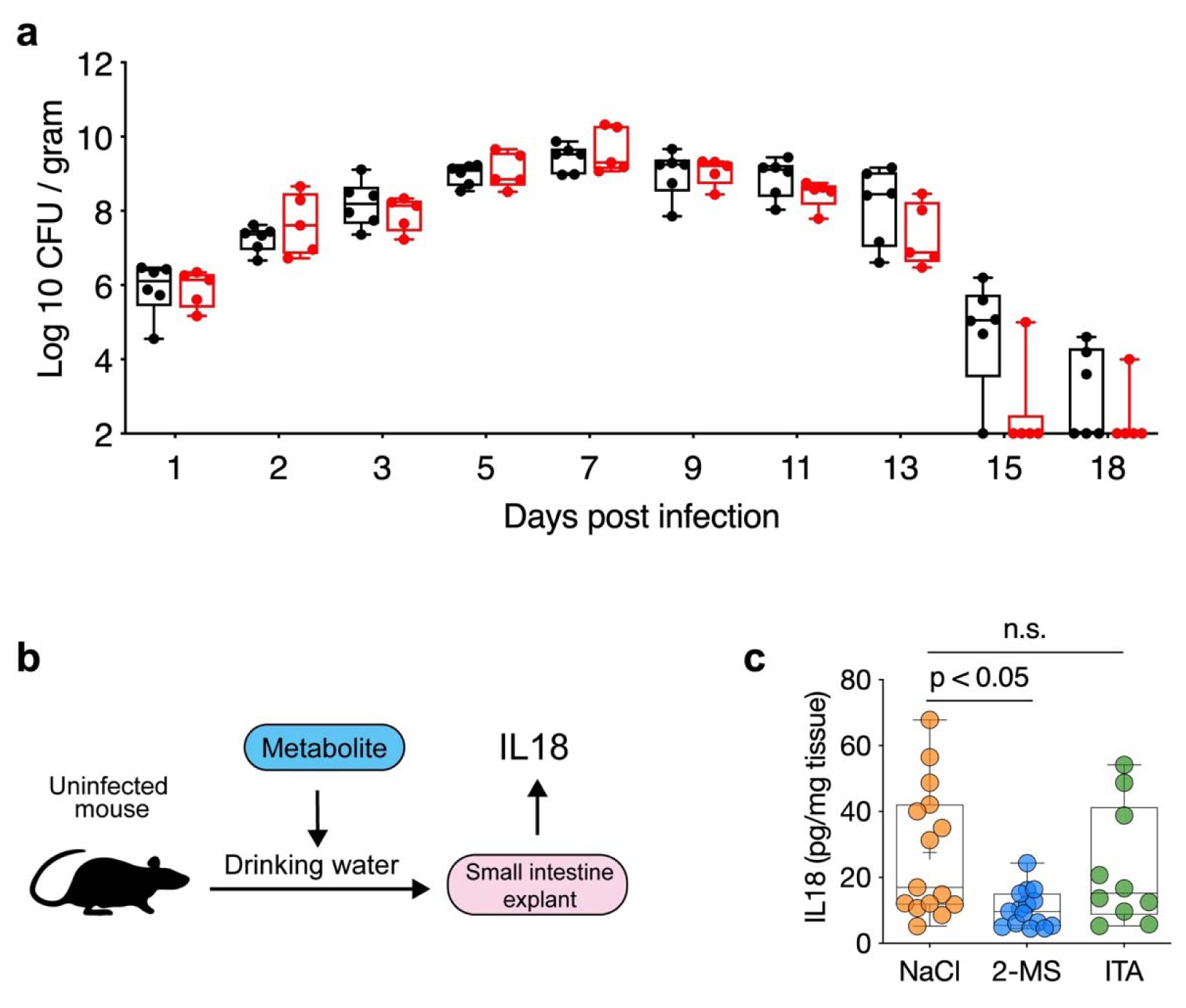
2-methylsuccinate suppresses intestinal inflammasome activity. (a) CFU of *C. rodentium* in the feces of infected WT and Acod1^-/-^ mice. (b) Experimental scheme for studying the effects of dicarboxylates on inflammasome activation. 5mM of individual dicarboxylate (adjusted to pH7 with NaOH) were placed into the drinking water of uninfected mice for 8 days, sodium concentration matched NaCl was used as control. Duodenal sections were dissected and cultured *ex vivo* for 20h as described ^60^. The abundance of IL-18 in supernatants was measured by ELISA. (c) Abundance of IL18 in supernatants of indicated duodenal explant cultures. Data are from 3 independent experiments with 5 mice/group for the NaCl treated control group and 2-methylsuccinate treated group, and 2 independent experiments in the itaconate treated groups.

**Supplementary Figure 8.**
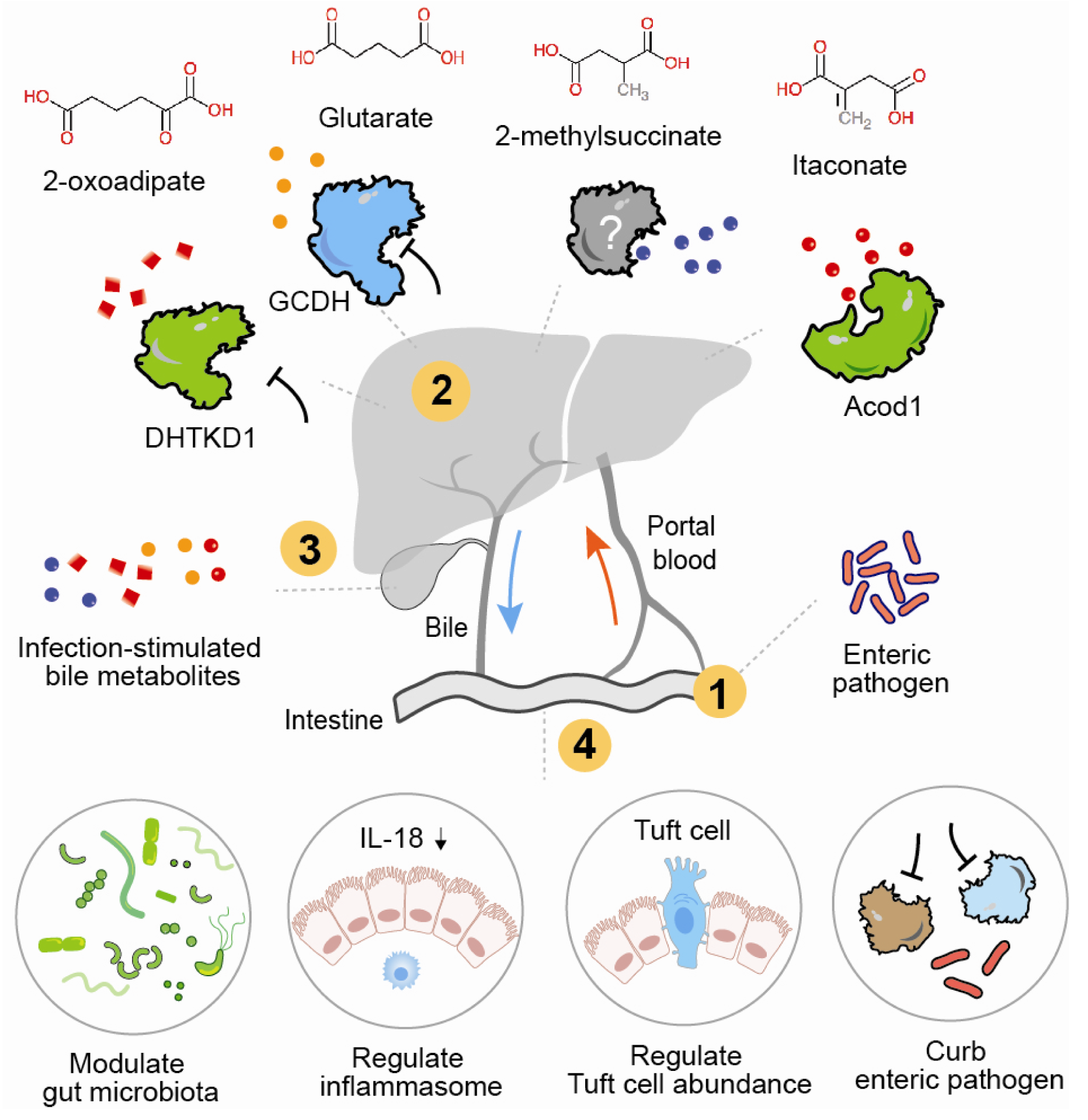
Bile remodeling triggered by infection modifies intestinal components and defense. Proposed model of a gut-liver defense circuit in which intestinal infection stimulates modifications in bile composition that in turn modulate intestinal function. Enteric infection (1) stimulates changes in hepatic transcriptional profiles (2) that alter bile composition (3) that in turn control microbiota and epithelial composition, inflammasome activity, and host defense (4).

